# Age-dependent deterioration of nuclear pore assembly in mitotic cells decreases transport dynamics

**DOI:** 10.1101/477802

**Authors:** I.L. Rempel, M.M. Crane, A. Mishra, D.P.M. Jansen, G.E. Janssens, P. Popken, M. Kaeberlein, E. Van der Giessen, P.R. Onck, A. Steen, L.M. Veenhoff

## Abstract

Nuclear transport is facilitated by the Nuclear Pore Complex (NPC) and is essential for life in eukaryotes. The NPC is a long-lived and exceptionally large structure. We asked whether NPC function is compromised in ageing mitotic cells. By imaging of single yeast cells during ageing, we show that the abundance of several NPC components and NPC assembly factors decreases while signs of misassembled NPCs appear. Consequently, nuclear permeability decreases, resulting in decreased dynamics of transcription factor shuttling and increased nuclear compartmentalisation. In support that declining NPC quality control is important in mitotic ageing, we find that the transport kinetics observed in ageing is mimicked in an NPC assembly mutant. Additionally, the single cell life histories reveal that cells that better maintain NPC function are longer lived. We conclude that assembly and quality control of NPCs are major challenges for ageing mitotic cells.

## Introduction

Rapid and controlled transport and communication between the nucleus and cytosol are essential for life in eukaryotes and malfunction is linked to cancer and neurodegeneration (reviewed in Fichtman and Harel, 2014). Nucleocytoplasmic transport, is exclusively performed by the Nuclear Pore Complex (NPC) and several nuclear transport receptors (NTRs or karyopherins) (reviewed in (Fiserova and Goldberg, 2010; Hurt and Beck, 2015)). NPCs are large (~30 MDa in yeast and ~50 MDa in humans) and dynamic structures(Alber et al., 2007; Kim et al., 2018; Onischenko et al., 2017; Teimer et al., 2017). Each NPC is composed of ~30 different proteins, called nucleoporins or Nups (Fig. 1a). The components of the symmetric core scaffold are long lived both in dividing yeast cells and in postmitotic cells, while several FG-Nups are turned over (D’Angelo et al., 2009; Denoth-Lippuner et al., 2014; Savas et al., 2012; Thayer et al.; Toyama et al., 2013) and dynamically associate with the NPC (Dilworth et al., 2001; Niño et al., 2016; Rabut et al., 2004). Previous studies performed in post-mitotic (chronologically ageing cells) showed changes in NPC structure and function (D’Angelo et al., 2009), but the fate of NPCs in ageing mitotic cells (replicative ageing) is largely unknown. To study the fate of NPCs in mitotic ageing we use replicative ageing budding yeasts as a model. Individual yeast cells have a finite lifespan defined as the number of divisions that they can go through before they die, their replicative lifespan (reviewed in (Longo et al., 2012)) (Fig. 1b). The divisions are asymmetric and while the mother cell ages the daughter cells are born young. Remarkably, studying the lifespan of this single cell eukaryote has been paramount for our understanding of ageing (reviewed in (Denoth Lippuner et al., 2014; Longo et al., 2012; Nyström and Liu, 2014)) and many of the changes that characterize ageing in yeast are shared with humans (Janssens and Veenhoff, 2016a). In the current study, we address changes to the NPC structure and function during mitotic ageing by imaging of single-cells.

**Fig. 1:**
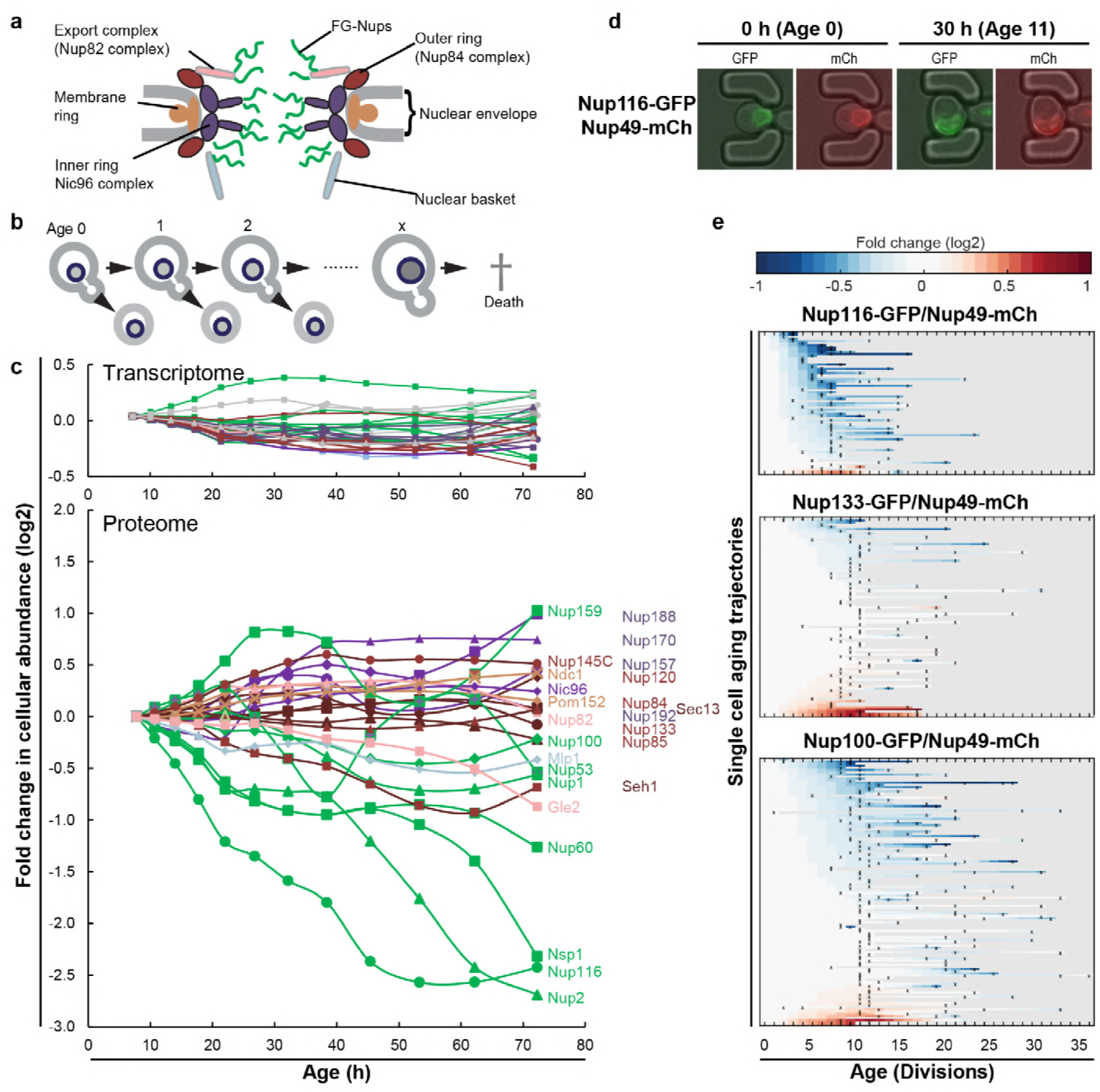
The cellular abundance of some NPC components changes in replicative ageing. **a,** Cartoon representation of the NPC (adapted from Kim et al., 2018) illustrates different structural regions of the NPC, all FG-Nups are shown in green independently of their localization, the membrane rings in light brown, the inner rings in purple, the outer rings in brown, the mRNA export complex in pink, and the nuclear basket structure in light blue. **b,** Schematic presentation of replicative ageing yeast cells. **c,** Transcript and protein abundance of NPC components (colour coded as in Fig. 1a) as measured in whole cell extracts of yeast cells of increasing replicative age; after 68 hours of cultivation the average replicative age of the cells is 24. Cells were aged under controlled and constant conditions. Data from Janssens et al., 2015. See also Supplementary Fig. 1a. **d,** Young cells are trapped in the microfluidic device and bright field images are taken every 20 minutes to define the cells age and fluorescent images are taken once every 15 hours to detect the protein localization and abundance. Representative images of cells expressing indicated fluorescent protein fusions imaged at the start of the experiment and after 30 hours; their replicative age is indicated. Scale bar represents 5µm. **e,** Heat map representation of the changes in the levels of the indicated GFP-and mCh-tagged Nups at the NE in each yeast cell at increasing age. Each line represents a single cell’s life history showing the change in the ratio of the fluorescence from the GFP-tagged Nup over the fluorescence of the mCh-tagged Nup and normalized to their ratio at time zero. Measurement of the fluorescence ratios are marked with “x”; in between two measurements the data was linearly interpolated. The fold changes are color coded on a log 2 scale from -1 to +1; blue colors indicate decreasing levels of the GFP-fusion relative to mCh. Number of cells in the heatmaps are Nup116-GFP/Nup49-mCh = 67, Nup133-GFP/Nup49-mCh = 94 and Nup100-GFP/Nup49-mCh = 126.

## Results

### The cellular abundance of specific NPC components changes in replicative ageing

We previously generated the first comprehensive dynamic proteome and transcriptome map during the replicative lifespan of yeast (Janssens et al., 2015), and identified the NPC as one of the complexes of which the stoichiometry of its components changes strongly with ageing. Indeed, the proteome and transcriptome give a comprehensive image of the cellular abundance of NPC components in ageing (Fig. 1c). We observe that the cellular levels of NPC components showed loss of stoichiometry during replicative ageing, which were not reflected in the more stable transcriptome data (Fig. 1c; Supplementary Fig. 1a). Clearly in mitotic ageing a posttranscriptional drift of Nup levels is apparent.

The total abundance of NPC components measured in these whole cell extracts potentially reflects an average of proteins originating from functional NPCs, prepores, misassembled NPCs, and possibly protein aggregates. Therefore, we validated for a subset of Nups (Nup133, Nup49, Nup100, Nup116 and Nup2) that GFP-tagged proteins expressed from their native promotors still localized at the nuclear envelope in old cells. In addition, we validated that changes in relative abundance of the Nups at the nuclear envelope were in line with the changes found in the proteome. We used microfluidic platforms that allow uninterrupted life-long imaging of cells under perfectly controlled constant conditions(Crane et al., 2014) (Fig. 1d). The single cell data of cells expressing GFP-fusions of Nup133 and Nup116 together with mCherry (mCh) fusions of Nup49 are shown in Fig. 1e (see Supplementary Fig. 2a-h for the other Nups and technical controls). Consistent with the proteome data, and with previously reported data (Lord et al., 2015), in the vast majority of ageing cells the abundance of Nup116-GFP decreased relative to Nup49-mCh, while the abundance of Nup133-GFP appears more stable. Also for the other Nups tested, the imaging data align well the proteome data (Supplementary Fig. 2c,f).

Our data contains full life histories of individual cells and, in line with previous reports (Crane et al., 2014; Fehrmann et al., 2013; Janssens and Veenhoff, 2016b; Jo et al.; Lee et al.; Zhang et al., 2012), we observed a significant cell to cell variation in the lifespan of individual cells, as well as variability in the levels of fluorescent tagged proteins. Therefore, we could assess if the changes observed for the individual NPC components correlated to the lifespan of a cell and, indeed, for Nup116 and Nup100 such correlations to lifespan were found, where those cells with lowest levels of NE-localized GFP-tagged Nups had the shortest remaining lifespan (for Nup100 r=0.48; p=1.27×10^-7^ and Nup116 r=0.56; p=6.54×10^-4^, see Supplementary Fig. 2g,h). The statistics of these correlations are in line with ageing being a multifactorial process where the predictive power of individual features is limited. In comparison to the ageing related increase in cell size (a Pearson correlation of around 0.2)(Janssens and Veenhoff, 2016b), the correlations found here are relatively large.

Taken together, we confirmed the loss of specific FG-Nups by quantifying the localization and abundance of fluorescently-tagged Nups in individual cells during their entire lifespan. Single cell Nup abundances at the NE can be highly variable (Nup2), while for other Nups (Nup100, Nup116 and Nup49) the loss in abundance at the NE was found in almost all ageing cells and correlated with the lifespan of the cell (Nup100 and Nup116). From the joint experiments published by Janssens et al., (Janssens et al., 2015); Lord et al., (Lord et al., 2015) and the current study we can conclude that especially Nup116 and Nsp1 (Nup98 and Nup62 in humans) strongly decrease in ageing.

### Mitotic ageing is associated with problems in NPC assembly rather than oxidative damage

A possible cause for the loss of stoichiometry could be that NPCs are not well maintained in ageing. Indeed, in postmitotic cells oxidative damage was proposed to lead to the appearance of carbonyl groups on Nups inducing more permeable NPCs (D’Angelo et al., 2009). We have limited information on the maintenance of existing NPCs during replicative ageing but there is some precedent for the hypothesis that even in the fast dividing yeast cells damage to existing NPCs may accumulate in aged cells. Indeed NPCs remain intact during multiple divisions (Colombi et al., 2013; Denoth-Lippuner et al., 2014; Khmelinskii et al., 2012; Thayer et al.), and especially in aged mother cells a fraction of the NPCs inherits asymmetrically to the ageing mother cell (Denoth-Lippuner et al., 2014). Oxidative stress and Reactive Oxygen Species (ROS) production in the cell is a major source of damage and can result in irreversible carbonylation of proteins (Stadtman and Levine, 2003). Protein carbonyls can be formed through several pathways. Here, we focused on the most prominent one, the direct oxidation of the

Lysine, Threonine, Arginine and Proline (K, T, R, P) side chains through Metal Catalyzed Oxidation (MCO)(Stadtman and Levine, 2003) by the Fenton reaction (Maisonneuve et al., 2009; Stadtman and Levine, 2003). Despite extensive efforts and using different in vitro and in vivo oxidative conditions and using different carbonyl-detection methods we could not find evidence for oxidative damage on several of the FG-Nups (Supplementary Fig. 4a,b showing negative results for Nsp1 along with a positive control).

Further indication that oxidative damage is unlikely to impact the NPC in ageing came from modelling studies. We carried out coarse-grained molecular dynamics simulations using our previously developed one-bead-per-amino-acid model of the disordered phase of the NPC (Ghavami et al., 2013, 2014). Earlier studies have shown that this model faithfully predicts the Stokes radii for a range of FG-domains/segments (Ghavami et al., 2014; Yamada et al., 2010), as well as the NPC’s size-dependent permeability barrier (Popken et al., 2015). To model the carbonylated FG-Nups, we incorporated the change in hydrophobicity and charge for carbonylated amino-acids (T, K, R, P) into the coarse-grained force fields (see Methods) and modelled maximally carbonyl-modified FG-Nups and NPCs. Overall, there is a minor impact of carbonylation on the predicted Stokes radius of the individual Nups and the time-averaged density of a wild type and fully oxidized NPC, with average densities around 80 mg/ml and maximum densities reaching 100 mg/ml in the center of the NPC (*r* < 5 nm) (Fig. 2a red line, Fig. 2b right panel and see Supplementary Fig. 4c-e for individual Nups and additional models dissecting the relative impact of the change in charge and hydrophobicity upon carbonylation).

**Fig. 2:**
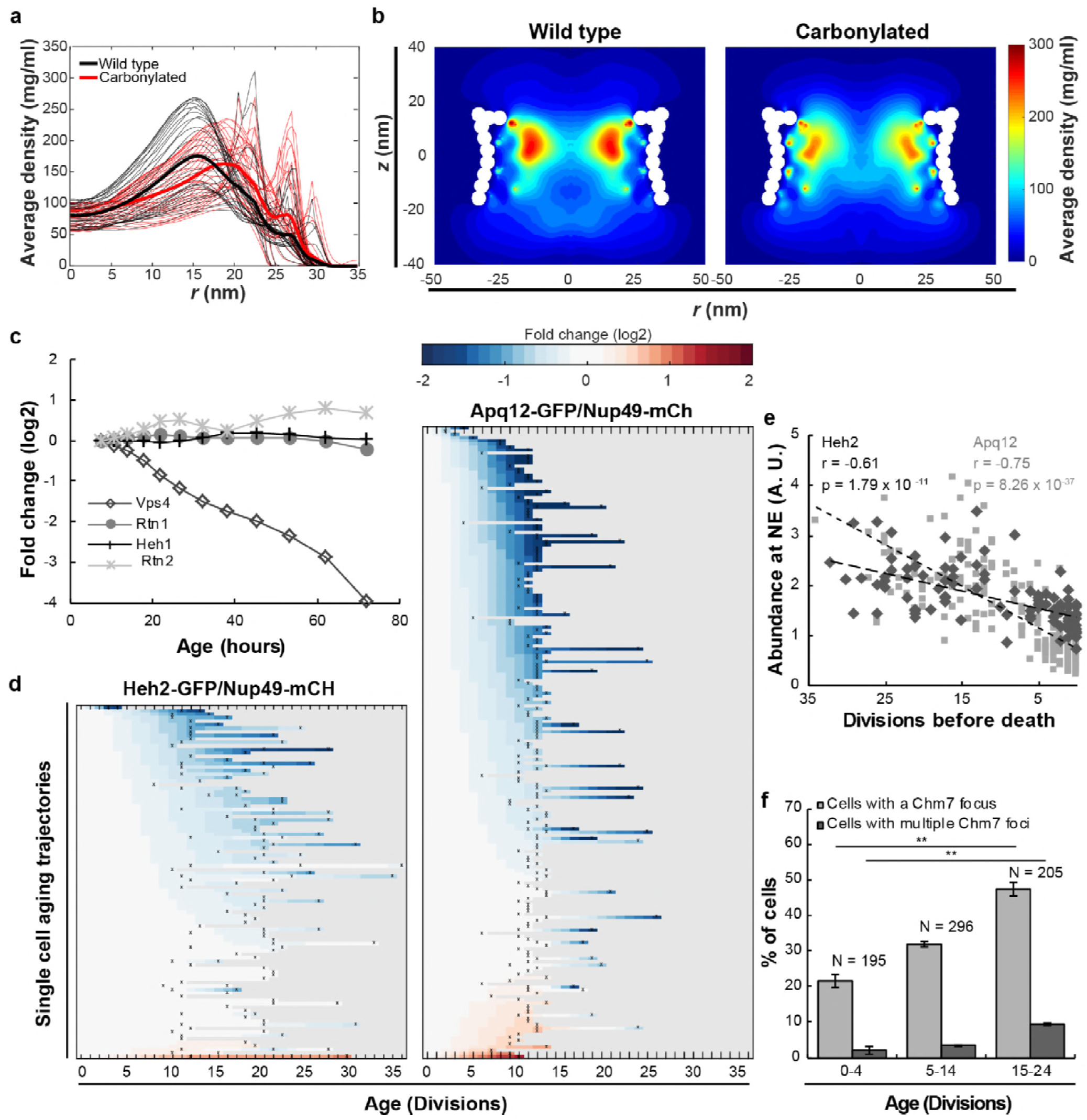
Mitotic ageing is associated with problems in NPC assembly rather than oxidative damage. **a,** Time-averaged radial density distribution of FG-Nups for different positions along the *z*-axis separated by 1 nm, in the range -15.4 < *z* < 15.4 nm, plotted for the wild type (black), the maximally carbonylated NPC (red) (See also Supplementary Fig. 4d,e). The dark coloured lines represent the density averaged over the range -15.4 < *z* < 15.4 nm. **b,** Time-averaged *r-z* density of FG-Nups in the wild type NPC (left panel), the oxidized NPC (right panel). **c,** Protein abundance of Heh1, Vps4, Rtn1 and Rtn2 as measured in whole cell extracts of yeast cells of increasing replicative age. Data from (Janssens et al., 2015). **d,** Heatmaps showing single cell abundance of Heh2-GFP (N = 100) and Apq12 (N = 200) at the NE, relative to Nup49-mCh in replicative ageing. **e,** Heh2-GFP and Apq12-GFP abundance at the NE, relative to Nup49-mCh, as a function of remaining lifespan. The dotted lines indicate best linear fit; Pearson correlations are indicated. Number of cells analyzed are Apq12 = 82, Heh2 = 51 and number of measuring points analyzed are Apq12 = 193 and Heh2 = 102. Data represents single replicates, a second replicate is shown in Supplementary Fig. 3. **f**, Percentage of cells with a Chm7 focus at the NE at different ages. Buds were excluded from the analysis. Error bars are weighted SD from the mean, from three independent replicates. P-values from Student’s t-test **p≤0.01. N = Total number of cells.

Altogether, we find no experimental evidence for carbonyl modification of FG-Nups even under strong oxidative conditions and, based on our modelling studies the carbonylation of FG-Nups is predicted to have little, or no impact on the passive permeability of NPCs, even under the unrealistic condition that all FG-Nups are fully carbonylated. We conclude that oxidative damage is unlikely to be a direct cause of altered NPC stoichiometry in replicative ageing, and it is probable that the previously reported increase in permeability of NEs during chronological ageing (D’Angelo et al., 2009) is actually caused by factors other than carbonylation.

We then addressed, if a main driver of NPC decline in replicative ageing may be the inability to control assembly. In young and healthy yeast cells misassembled NPCs are absent, but mutant strains with impaired NPC assembly show that a fraction of their NPCs cluster, are covered by membranes, or cause herniations of the NE (Chadrin et al., 2010; Scarcelli et al., 2007; Webster et al., 2014, 2016) (reviewed in Thaller and Lusk, 2018). Misassembled NPCs that are induced by mutations are asymmetrically retained, and accumulated in the mother cell over time (Webster et al., 2014). We thus asked, if replicatively aged cells start to progressively accumulate misassembled NPCs. Correct NPC assembly is assisted by several proteins that are temporarily associated with NPCs during the assembly process (Dawson et al., 2009; Lone et al., 2015; Otsuka and Ellenberg, 2018; Scarcelli et al., 2007; Webster et al., 2016; Zhang et al., 2018). Amongst these are (i) Heh1 and Heh2, the orthologues of human LEM2 and Man1, which are known to be involved in recognition of misassembled pores (Webster et al., 2014, 2016), (ii) Vps4, an AAA-ATPase with multiple functions amongst which the clearance of misassembled NPCs from the NE (Webster et al., 2014) and (iii) Apq12, Rtn1 and Rtn2, Brr6 and Brl1 membrane proteins of the NE-ER network that are involved in NPC assembly, possibly through roles in modulating membrane curvature (Lone et al., 2015; Scarcelli et al., 2007; Zhang et al., 2018).

The system wide proteomics data showed that the protein levels of Heh1, Rtn1 and Rtn2 are stable in abundance in ageing, while a sharp decrease in abundance was found for Vps4 (Fig. 2c, and Supplementary Fig. 1c showing stable transcript levels). Additionally, we found that the abundance of Heh2-GFP and Apq12-GFP at the NE decreased relative to Nup49-mCh in ageing (Fig. 2d, and Supplementary Fig. 1c showing stable transcript levels). Despite the fact that neither Heh2 nor Apq12 are essential proteins, we found their levels to be correlated with the remaining lifespan of the cells, where those cells showing the lowest levels of Heh2-GFP or Apq12-GFP had the shortest remaining lifespan (Fig. 2e). Previous work showed that a single deletion of *heh2*, *vps4* or *apq12* is sufficient to cause misassembled NPCs (Scarcelli et al., 2007; Webster et al., 2014) so the decrease in abundance of the proteins Heh2, Apq12 and Vps4 suggests that NPC assembly is compromised in ageing and misassembled NPCs may accumulate.

To get a more direct readout of problems in NPC assembly we studied Chm7, the nuclear adaptor for the ESCRT system (Webster et al., 2016). Chm7 sometimes forms a focus at the NE and the frequency of focus formation is related to NPC assembly problems as mutant strains with impaired NPC assembly show more frequently Chm7 foci at the NE (Webster et al., 2016). We quantified the frequency of focus formation in differently aged cells. Indeed, the foci are more than twice as frequently seen in the highest age group (age 15-24), compared to cells younger than 5 divisions. Also, the frequency at which cells have more than one focus present at the NE is more than 4-fold higher in the oldest age group (Fig. 2f).

We conclude that three proteins involved in the assembly of NPCs decrease strongly in abundance in ageing (Vps4, Heh2 and Apq12) in a manner that correlates with remaining lifespan (Fig. 2). The decrease in abundance of those proteins, and potentially also the decrease of FG-Nup abundance (Fig. 1), is likely to cause the NPC assembly problems, which we observe as an increased Chm7 focus formation frequency (Fig. 2f).

### Increased steady state nuclear compartmentalization in ageing is mimicked in an NPC assembly mutant

Next, we experimentally addressed the rates of transport through and from the nucleus with ageing. During import and export NTRs bind their cargoes through a nuclear localization signal (NLS) or nuclear export signal (NES) and shuttle them through the NPC. In addition to facilitating active transport, the NPC is a size dependent diffusion barrier (Popken et al., 2015; Timney et al., 2016). We measured the rate of efflux in single ageing cells and find that passive permeability is not altered significantly in ageing (Supplementary Fig. 5a-c), excluding the possibility that that NPCs with compromised permeability barriers (‘leaky’ NPCs) are prevalent in ageing cells. We then looked at classical import facilitated by the importins Kap60 and Kap95, and export facilitated by the exportin Crm1. The cellular abundance of Crm1, Kap60 and Kap95 is relatively stable in ageing (Janssens et al., 2015) (Supplementary Fig. 6a and Supplementary Fig. 1c for transcript levels) as is their abundance at the NE and their localisation (Supplementary Fig. 6b-d). To test whether transport kinetics change with ageing, we used GFP-NLS (Kap60, Kap95 import cargo) and GFP-NES (Crm1 export cargo) reporter proteins, and GFP as a control. The steady state localization of these proteins depends primarily on the kinetics of NTR facilitated transport (import or export) and passive permeability (influx and efflux), while retention mechanisms are minimal, enabling direct assessment of transport kinetics.

We carefully quantified the steady state localisation of transport reporters in individual ageing cells in the non-invasive microfluidic setup (See Supplementary Fig. 7 for lifespan of strains). In the vast majority of cells we observed that GFP carrying a NLS accumulated more strongly in the nucleus at high ages (Fig. 3a, middle panel), and, interestingly, the GFP carrying a NES is more strongly depleted from the nucleus in the vast majority of cells (Fig. 3a, right panel). For the control, GFP, we find a more stable N/C ratio in ageing (Fig. 3a, left panel). While the changes in steady state accumulation are observed already early in life when looking at single cells, on the population level the changes become significant only later in the lifespan (Fig. 3b). A previous report showed a reduction in compartmentalization of GFP-NLS in age 6+ yeast cells isolated from a culture (Lord et al., 2015), while we see no statistically significant difference at this age. We note that there are many differences in the experimental setups that may explain the difference. The systematic changes in the steady state localisation of the reporter proteins that we observe in the ageing cells show that in aged cells the balance between the rates of NTR-facilitated-transport (import and export) and passive permeability (influx and efflux) are altered such that in old cells the kinetics of passive permeability is lowered relative to the kinetics of NTR-facilitated-transport.

**Fig. 3:**
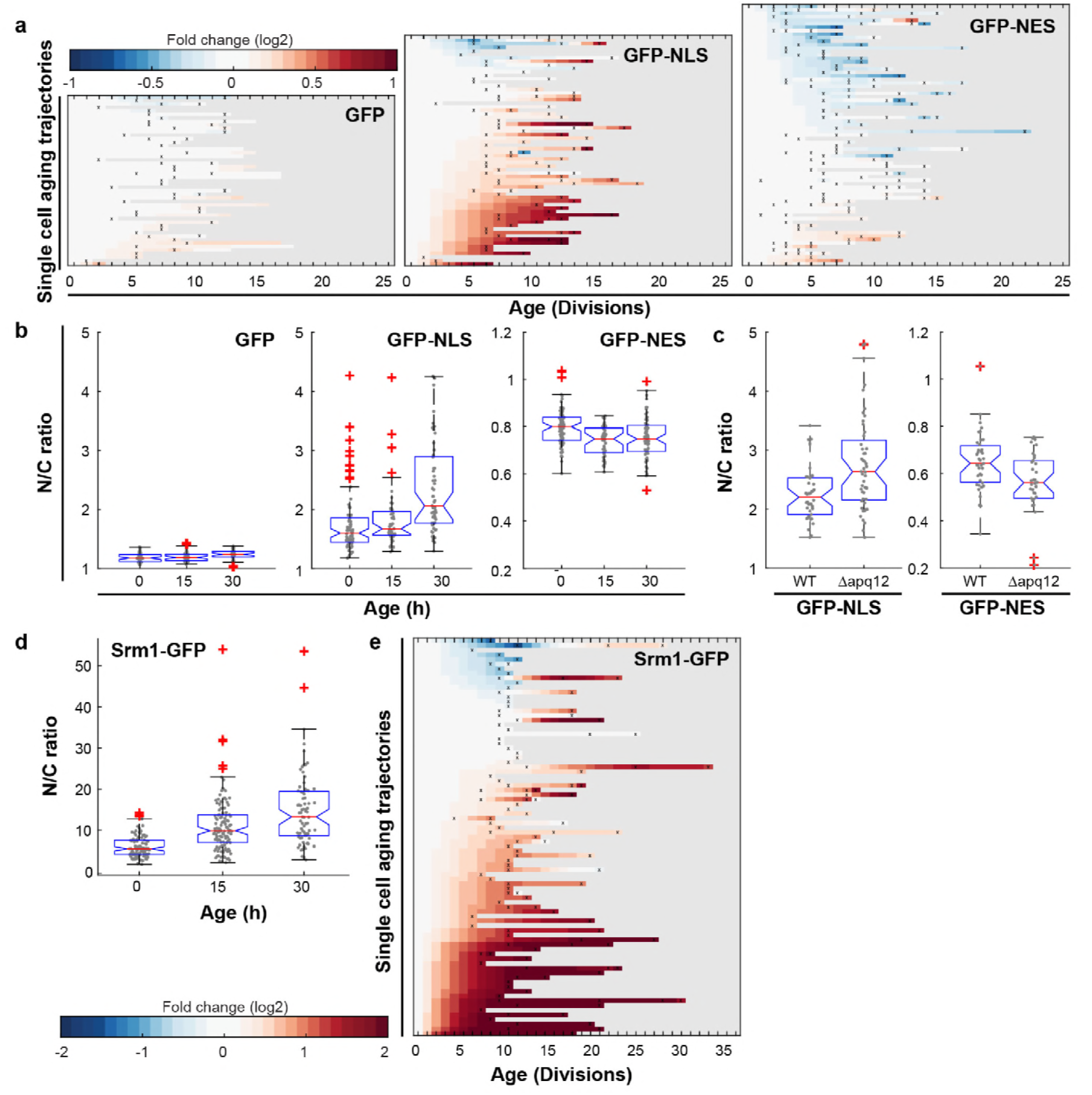
Increased steady state nuclear compartmentalization in ageing is mimicked in an NPC assembly mutant. **a,** Heatmaps showing single cell changes in localization (N/C ratios) of GFP (N = 49), GFP-NES (N = 75) and GFP-NLS (N = 66) reporter proteins during replicative ageing. **b,** N/C ratios of GFP-NLS, GFP-NES and GFP as the cells age. The line indicates the median, and the bottom and top edges of the box indicate the 25th and 75th percentiles, respectively. The whiskers extend to the most extreme data points not considered outliers, and the outliers are plotted individually. Non overlapping notches indicate that the samples are different with 95% confidence. The number of cells analyzed are GFP = 54, 51, 34; GFP-NLS = 74, 48, 57 and GFP-NES = 75, 41, 66 after 0, 15 and 30 hours respectively. **c,** Deletion of *apq12* increases nuclear compartmentalization of GFP-NLS and GFP-NES. The number of cells analyzed are GFP-NLS = 42, 48 and GFP-NES = 39, 34 for WT and Δapq12 respectively **d,** N/C ratios of Srm1-GFP increases as cells age. Numbers of cells analyzed are N = 103, 125, 77 after 0, 15 and 30 hours respectively **e,** Heatmap showing single cell changes in localization (N/C ratios) of Srm1-GFP (N = 85) during replicative ageing.

To our knowledge, there are no strains with mutations in the NPC that have been shown to lead to increased steady state compartmentalisation. On the contrary, many strains, including those where NPC components that decrease in abundance in ageing are deleted or truncated, show loss of compartmentalisation (Lord et al., 2015; Popken et al., 2015; Strawn et al., 2004). Interestingly, the only strain that was previously reported to have an increased compartmentalisation is a strain defective in NPC assembly due to a deletion of *apq12* (Scarcelli et al., 2007; Webster et al., 2016). We found that deletion of *apq12* is not viable in the BY strain background, hence we recreated the deletion mutant in the W303 background, where it is stable. Indeed, we found that the deletion of *apq12* was sufficient to mimic the increase in compartmentalization seen in ageing showing increased nuclear accumulation of GFP-NLS and exclusion of GFP-NES (Fig. 3c). The increased compartmentalisation in aged cells and in the *apq12* mutant can be explained if fewer functional NPCs are present in the NE. Reduced numbers of NPCs would predominantly impact passive permeability, as the rate limiting step for NTR-facilitated-transport is not at the level of the number of NPCs but rather at the level of NTRs and cargo’s finding each other in the crowded cytosol with overwhelming nonspecific competition (Meinema et al., 2013; Riddick and Macara, 2005; Smith et al., 2002; Timney et al., 2016).

Next, we addressed how the altered transport dynamics in aged cells impacts native proteins. We studied Srm1, the yeast homologue of Rcc1, as endogenously expressed GFP-tagged protein. Srm1 is the nucleotide exchange factor that exchanges GDP for GTP on Ran and its nuclear localization ensures that Ran-GTP levels inside the nucleus are high. The localization of Srm1 depends on Kap60/Kap95 mediated import and retention inside the nucleus via chromatin binding (Li et al., 2003; Nemergut et al., 2001). While the cellular abundance of Srm1 was stable during ageing (Supplementary Fig. 1b), we found that the nuclear accumulation of Srm1-GFP increased during replicative ageing in most cells (Fig. 3d,e). The change in localization of Srm1-GFP is in line with the changes observed for GFP-NLS. Interestingly, the human homologue of Srm1, Rcc1, was previously also reported to have an increased nuclear concentration in myonuclei and brain nuclei of aged mice (Cutler et al., 2017).

### Alterations nuclear envelope permeability during ageing affects transcription factor dynamics

Additionally, we studied Msn2, a transcriptional regulator that responds to various stresses and translocates to the nucleus in pulses, a so called frequency modulated transcription factor. Msn2 communicates information about the environment by modifying the frequency of pulses (Fig. 4a)(Cai et al., 2008; Hao and O’Shea, 2011; Hao et al., 2013). These pulse dynamics are primarily determined by the rates of nuclear import and nuclear export (Hao et al., 2013). By following endogenously tagged Msn2-GFP, we were able to observe the pulse dynamics for individual cells, and quantify specific features of each pulse (Fig. 4b). Specifically, for each Msn2 pulse, we determined the peak prominence and the pulse width. To determine how ageing affected the import and export kinetics, we imaged mother cells for short periods of time every ten hours (Fig. 4c). We observed that, as predicted by the alterations in NPCs, the average pulse widths for each cell increased consistently from middle-age onwards (Fig. 4d). Similarly, by aligning all Msn2 pulses on top of each other for a given age, we determined a mean pulse shape for each age (Fig. 4e). These show similar characteristics, where the ageing results in both broader and lower Msn2 pulses. As could be expected given the striking age-related changes, both the pulse width and pulse prominence are correlated to the remaining lifespan of the cell (Supplementary Fig. 8a,b, r=-0.36, p<0.0001 and r=0.47, p<0.0001 for pulse prominence and pulse width respectively). We consider it likely, that the decrease in Msn2 dynamics is a readout for the NPC’s ability to facilitate rapid responses. Ageing cells may thus respond more slowly to changes in their environment due to reduced nuclear cytoplasmic transport and communication.

**Fig. 4:**
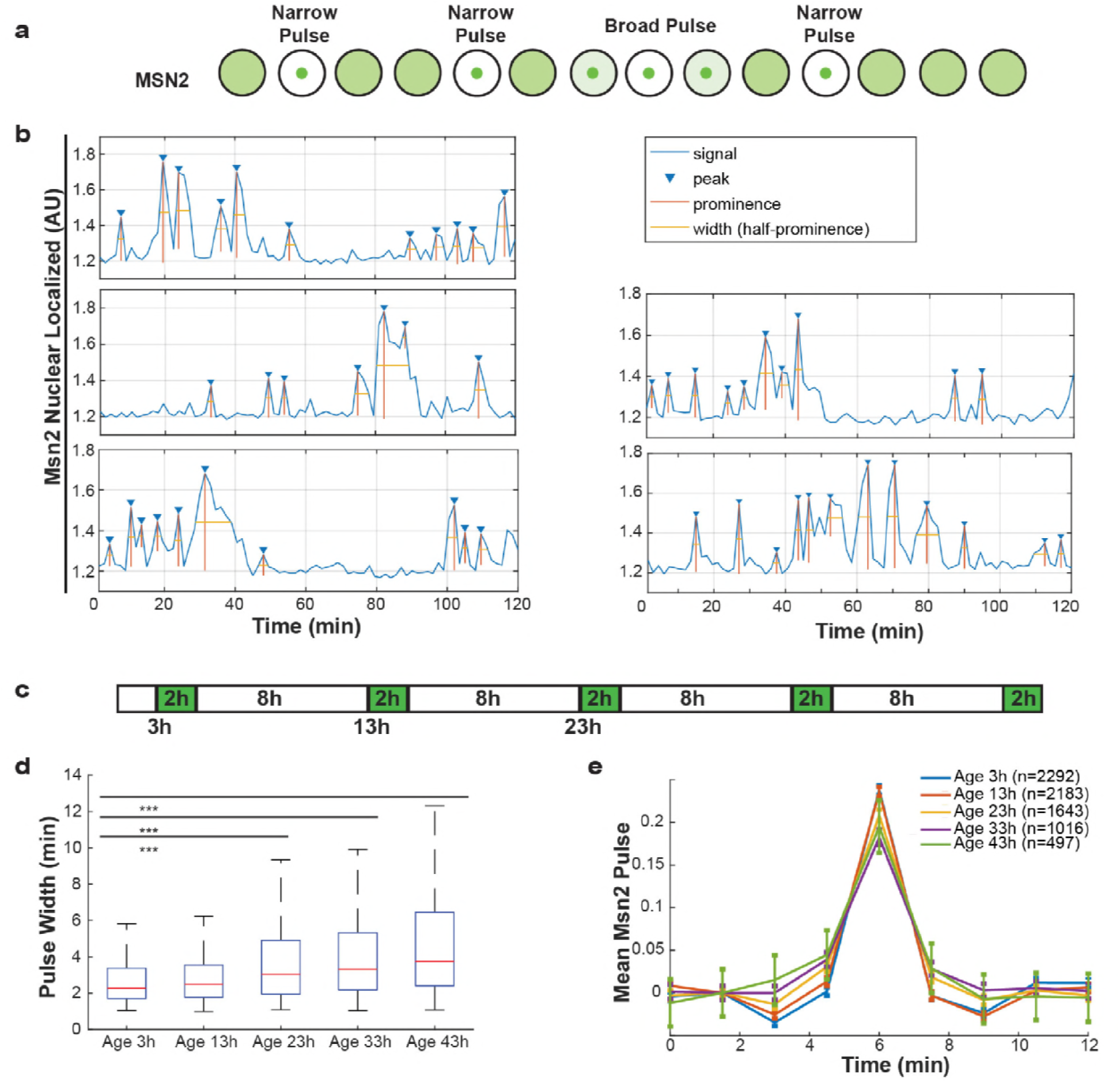
Alterations nuclear envelope permeability during ageing affects transcription factor dynamics. **a,** Schematic showing pulses of Msn2 translocation to the nucleus and movement back to the cytoplasm. **b,** Five randomly selected single cell traces showing Msn2 dynamics. Low values indicate the majority of Msn2 is cytoplasmic, and high values indicate the majority of Msn2 in nuclear localized. Pulses are annotated showing peaks, prominence and the width of the pulse. **c,** Experimental protocol for the ageing experiment. White boxes indicate brightfield imaging only, and green boxes indicate fluorescence imaging. **d,** As cells age, the width of the Msn2 pulses increases reliably. (*** indicates p<0.0001 two-tailed t-test). **e,** Msn2 pulses were identified at each age, and then all pulses were averaged together at each age. To correct for changes in baseline localization with age, the mean pre-pulse level was subtracted at each age. Error bars are standard error.

## Discussion

NPC function in ageing has received much attention in the context of chronological ageing cells such as neurons and indeed a large body of data now implicated NPC function in neurodegenerative diseases. Here we study the fate NPCs in dividing yeast cells with the anticipation that the insights may be relevant to ageing of mitotic cells such as stem cells. Overall our data is consistent with a model where NPC assembly and quality control is compromised in mitotic ageing (Fig. 2) and where misassembled NPCs that specifically lack the FG-Nups that we see declining in ageing (Fig. 1) accumulate in aged cells. Without intervention, the loss of FG-Nups at the NE would almost certainly create leaky NPCs (Popken et al., 2015; Strawn et al., 2004; Timney et al., 2016), as also predicted by our models of aged NPCs (Supplementary Fig. 9). However, based on our transport data, we can exclude a scenario where leaky NPCs are present at the NE of mitotically aged cells (Fig. 3,4). We consider it likely that the misassembled NPCs get covered with membrane, a structure known to be present in *nup116*, *vps4*, *heh2* and *apq12* mutant strains (Scarcelli et al., 2007; Webster et al., 2014, 2016; Wente and Blobel). This would prevent loss of compartmentalization but cause an overall reduction of permeable NPCs, which is consistent with the observed decrease in transport dynamics across the NE and the increased steady state compartmentalisation. As a result, aged nuclei thus have fewer functional NPCs and significant amounts of dysfunctional NPCs that do not contribute to the overall transport kinetics.

Mitotically aged cells show an increased compartmentalisation; a phenotype that is clearly distinct from the loss of compartmentalisation by leaky NPCs observed in aged post-mitotic cells (D’Angelo et al., 2009). Our finding of correlations between remaining lifespan of a cell and the abundance of several Nups and assembly factors, as well as the transport dynamics of Msn2, supports that a potential causal relation between NPC function and lifespan. Mitotically ageing cells are expected to respond more slowly to changes in their environment due to reduced nuclear cytoplasmic transport and communication. Moreover, declining NPC function, itself the consequence of failing quality control, will result in aberrant transcription factor regulation and mRNA export, and, with that, loss of protein homeostasis. Altogether, our study contributes to the emerging view that NPC function is a factor of importance in ageing and age-related disease (Fichtman and Harel, 2014) impinging on the universal hallmarks of human ageing of intercellular communication and loss of protein homeostasis.

Our work also provides the first clear example that the assembly of large protein complexes is a major challenge in ageing of dividing cells. Indeed, there is data to support that loss of protein complex stoichiometry is a prominent and conserved phenotype in ageing (Janssens et al., 2015; Ori et al., 2015). We speculate that the assembly and quality control of many other large protein complexes, such as the proteasome and kinetochores, both becoming highly substoichiometric in ageing (Janssens et al., 2015; Ori et al., 2015), are compromised in mitotically ageing cells.

## Materials and Methods

### Strains

All *Saccharomyces cerevisiae* strains used in this study are listed Table I and were validated by sequencing. Experiments in this study were performed with BY4741 genetic backgrounds, except the deletion of *apq12*, which is instable in the BY4741 background. W303 apq12Δ was created by using the PCR-toolbox (Janke et al., 2004). The Nup116-GFPboundary MKY227 from Mattheyses et al., 2010 was converted from its W303 background to BY4741 background by crossing and tetrad dissection for in total 10 times with BY4741. All strains used in the aging experiments are plasmid free as plasmids are not maintained in aging cells. Cells were grown at 30°C, shaking at 200 RPM using Synthetic Complete medium supplemented with 2% Glucose or 2% D-raffinose, unless indicated otherwise. If applicable, expression of reporter proteins was induced with 0.5% Galactose. Cells were induced for 4-7 h prior to the start of an experiment. The proteomics data within this manuscript previously published in Janssens et al., 2015 is from YSBN6 grown in nitrogen base without amino acids supplemented with 2% glucose.

**Table 1:**
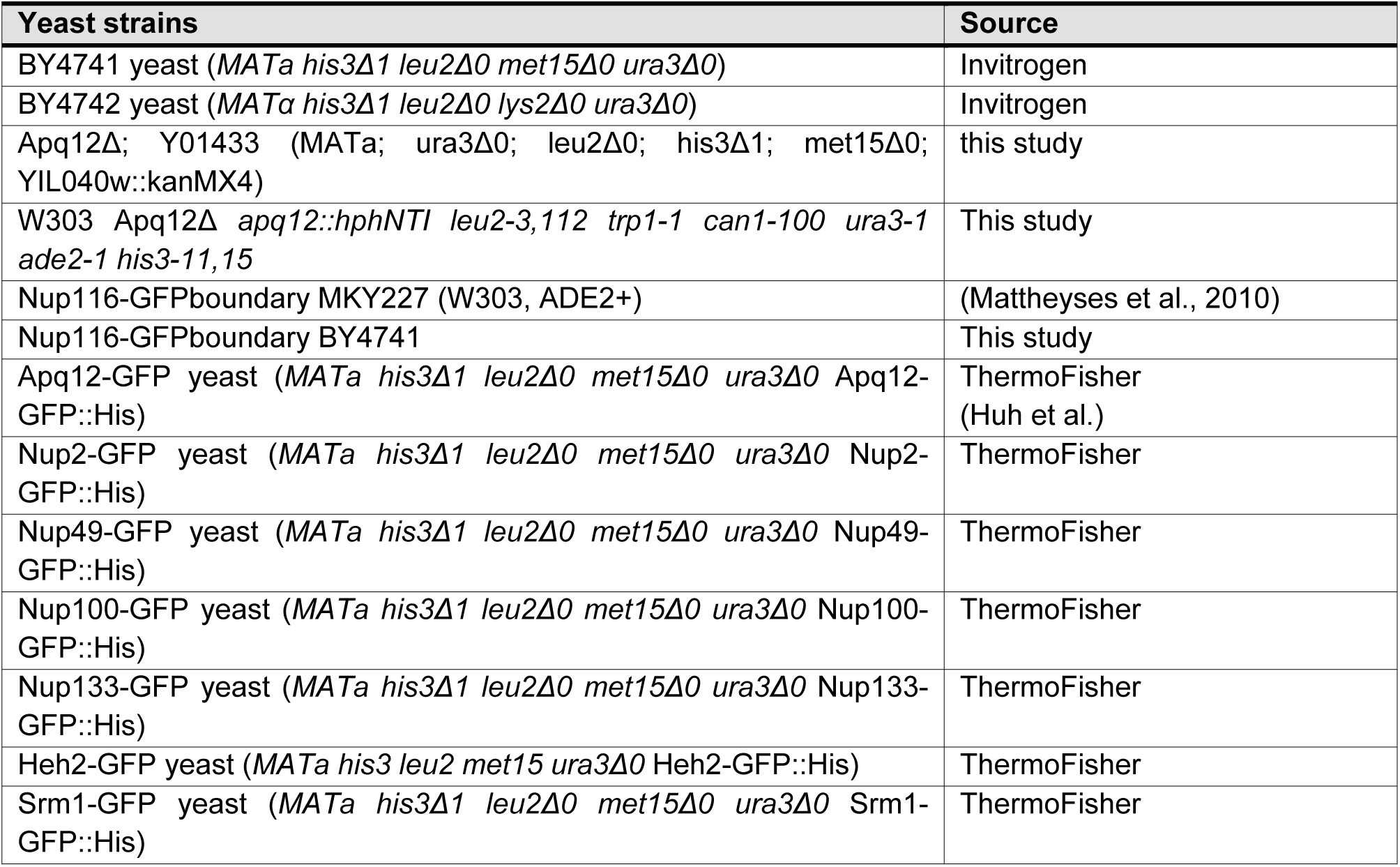
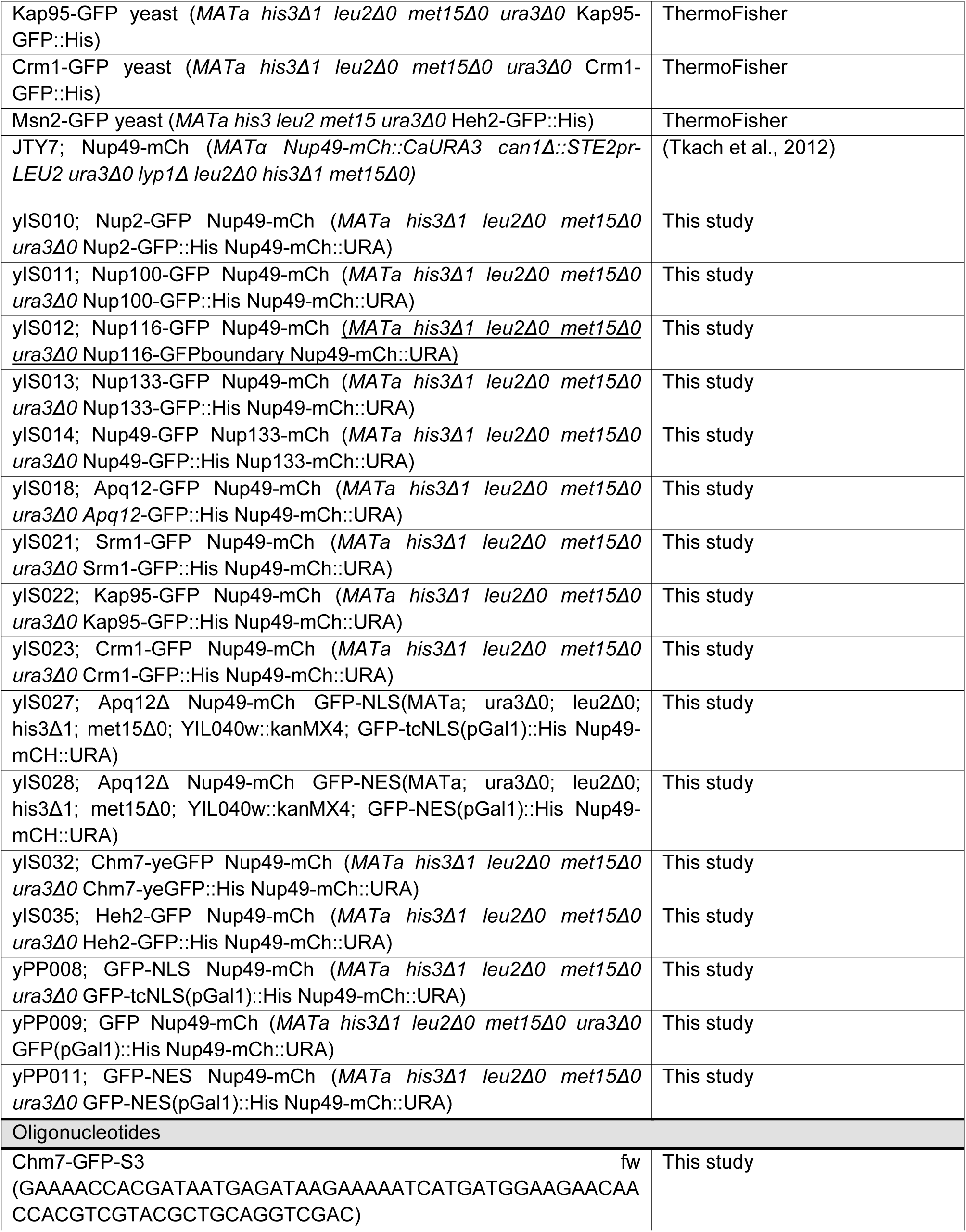
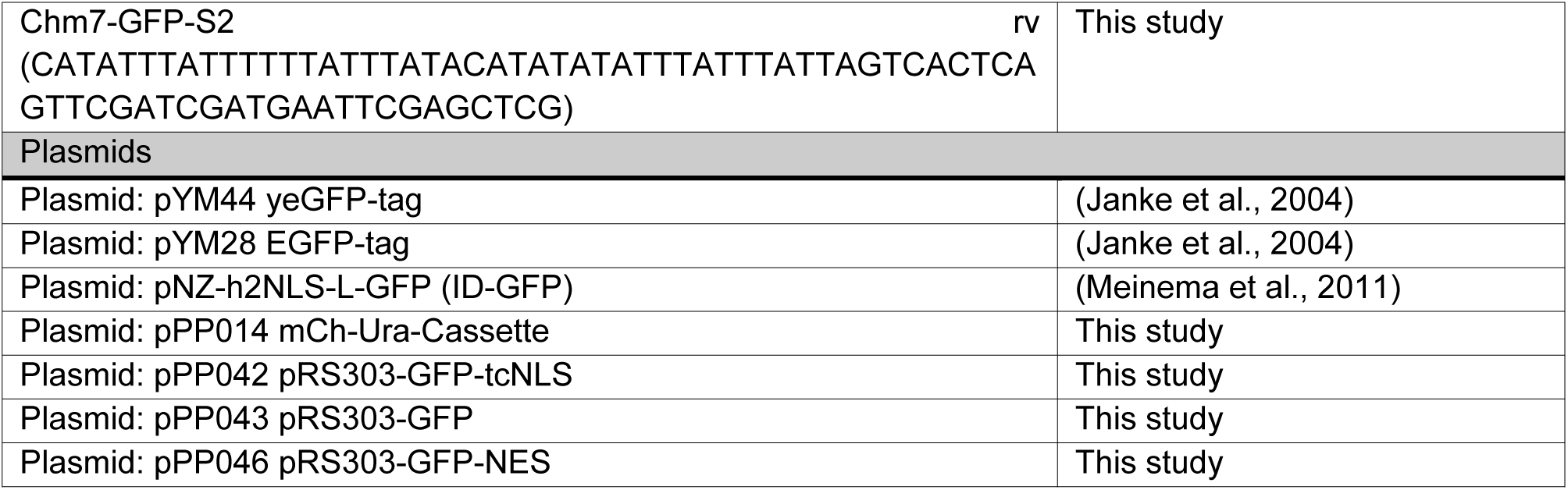
Strains and plasmids

### Microscopy

All microscopy, excluding the experiments for Fig. 7, was performed at 30°C on a Delta Vision Deconvolution Microscope (Applied Precision), using InsightSSITM Solid State Illumination of 488 and 594 nm, an Olympus UPLS Apo 60x or 100x oil objective with 1.4NA and softWoRx software (GE lifesciences). Detection was done with a CoolSNAP HQ2 camera. Microscopy to study Msn2 dynamics was performed on a Nikon Ti-E microscope equipped with a Hamamatsu Orca Flash V2 using a 40X oil immersion objective (1.3NA). Fluorescence excitation was performed using an LED illumination system (Excelitas 110-LED) that is triggered by the camera.

### Replicative aging experiments – microfluidic dissection platforms

The microfluidic devices were used as previously detailed (Crane et al., 2014; Lee et al.). Brightfield images of the cells were taken every 20 min to follow all divisions of each cell. Fluorescent images with three or four z-slices of 0.5 or 0.7 micron, were taken at the beginning of the experiment and after 15, 30, 45 and 60 h. One experiment lasts for a maximum of 80 h. All lifespans and the N/C ratios of yPP008, yPP009 and yPP011 (Figures 6a,c and d) reflect only cells that stay in the device for a whole lifespan are included into the dataset. All other data presented include all cells that stay in the microfluidic device for at least 15 h and have at least one image well enough in focus for a ratiometric measurement. Data in Fig. 6a,c and d was obtained using both the microfluidic dissection platform (Lee et al.) and the ALCATRAS (Crane et al., 2014), data in all other experiments were performed using an ALCATRAS chip.

### Poison assay in the microfluidic device

To measure the passive permeability of NPCs in old cells, the cells were replicatively aged in the microfluidic chip for approximately 21 h. Subsequently, the medium in the chip was exchanged, as described by (Crane et al., 2014), for Synthetic Complete medium supplemented with 10 mM sodium azide and 10 mM 2-deoxy-D-glucose (Shulga et al., 1996). Additionally, the medium was supplemented with some Ponceau S stain, which makes the medium fluoresce in the mCherry channel. The addition of sodium azide and 2-deoxyglucose depletes the cell of energy and destroyes the Ran-GTP/GDP gradient thus abolishing active transport of reporter proteins. We measured the net efflux of reporter proteins by imaging the cells every 30 s.

### Data analysis of Nups and N/C ratios

Microscopy data was quantified with open source software Fiji (https://imagej.net/Welcome)(Schindelin et al., 2012). Fluorescent intensity measurements were corrected for background fluorescence. To quantify the abundance of proteins at the NE, an outline was made along the NE in the mCherry channel. The outline was used to measure the average fluorescent intensities in the mCherry and GFP channels. To quantify the nuclear localization (N/C ratio), the NE was outlined based on the Nup49-mCherry signal and the average fluorescence intensity at the nucleus was measured. A section in the cytosol devoid of vacuoles (appearing black) was selected for determining the average fluorescence intensity in the cytosol. We note that the average fluorescence of GFP in the cytosol may be underestimated in aged cells as aged cells have many small vacuoles that make it hard to select vacuole-free areas in the cytosol. The extent to which this affects the data can best be judged from the cells expression GFP where the N/C ratio on average increases from 1.2 to 1.25 in 30 hours (Fig. 6d). All heatmaps and bee swarm/box plots were generated in MATLAB (Mathworks https://nl.mathworks.com/).

### Msn2:GFP imaging and quantification

Following introduction of the cells to the microfluidic device, brightfield imaging was begun immediately. The process of introducing cells to the device was found to increase Msn2 activity for the first hour or two following device loading. To ensure that the baseline timecourse in young cells was representative of pulse dynamics and not affected by loading stress, fluorescence imaging was begun after cells had been allowed to acclimate for three hours. Brightfield images were acquired at every timepoint, with intervals of five minutes when fluorescence images were not acquired. Fluorescence images were acquired for two hours, at intervals of 90 seconds, with three z-slices of 1.5 microns. Following the fluorescence imaging, brightfield images were acquired for 8 hours to ensure that cells could be tracked and the number of daughter divisions could be scored. Cells were segmented and tracked using previously published software (Bakker et al., 2018).

Nuclear accumulation of Msn2 was quantified using a measure of skewness. Specifically, the ratio of the brightest 2% of the pixels within the cell relative to the median cell fluorescence. By normalizing to the median cell fluorescence, this is measurement is robust to photobleaching or changes in protein concentration. This measurement has been repeatedly validated and used in previous studies of transcription factor translocation dynamics (Cai et al., 2008; Granados et al., 2017). For each single cell, peaks were located and quantified using the findpeaks function in Matlab®. At each age, the measurements for all pulses within a single cell were averaged to generate a single value for mean Msn2 pulse properties for that cell, at that age. This value was used for correlations with remaining lifespan and distribution of pulse widths at each age (Supplementary Fig. 8). To determine the mean pulse dynamics at each age, all pulses of all cells alive at the age were centered relative to each pulse peak, and averaged.

### Modelling of aged NPCs

In order to model the aged NPC with the measured stoichiometry of FG-Nups from the protein abundance data (see Fig. 1), we built 24 different models by taking into account the 8-fold symmetry of the NPC. The model details are shown in Table S1. In all 24 models, the two peripheral Nsp1’s along with Nup116 were deleted. Nsp1 in the central channel recruits Nup49 and Nup57 to form a Nsp1-Nup49-Nup57 subcomplex. As a result, deletion of one of the central channel Nsp1’s is accompanied by removal of the corresponding Nup49 and Nup57. We computed the time-averaged radial mass density distribution (density averaged in the circumferential and axial direction) of the FG-Nups for these 24 models along with the wild type (Supplementary Fig. 9). The average of the 24 models we refer to as the ‘Aged proteome’ model.

### Force-field parameters of carbonylated amino acids in the 1-bead-per-amino-acid (1BPA) model(Ghavami et al., 2014)

Among all amino-acids, Threonine (T), Lysine (K), Proline (P) and Arginine (R) can undergo carbonylation. The change in hydrophobicity upon carbonylation of these amino-acids was calculated with the help of five hydrophobicity prediction programs (KOWWIN, ClogP, ChemAxon, ALOGPS and miLogP) (Leo, 1993; Meylan and Howard, 1995; Tetko et al., 2005; Viswanadhan et al., 1989)(Cheminformatics, 2015, www.molinspiration.com). These software programs use the partition coefficient *P* (ratio of concentrations in a mixture of two immiscible phases at equilibrium, usually water and octanol) as a measure of hydrophobicity. Since the range of the ratio of concentrations is large, the logarithm of the ratio of concentrations is commonly used: 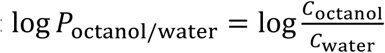.

These programs provide different estimates of the value of log *P* for a given chemical structure. They use experimental log*P* values for atoms or small groups of atoms as a basis and their algorithms are fine-tuned by training with experimental values of complete molecules. The molecules are cut into fragments or into atoms, and their contribution adds up to the log*P* value of the entire molecule based on the concept of structure-additivity (Fujita et al., 1964).

The hydrophobicity scale in the 1BPA force field(Ghavami et al., 2014) is derived from three scales that are based on partition energy measurements. Since the free energy of partition is proportional to log*P*, the strategy was chosen to find log*P* values for the oxidized amino acids. To obtain a reliable value for each of the chemically modified amino acids, a weighted average scheme is used. Instead of using the predicted hydrophobicity for the entire residue, it is more accurate to use the predicted change in hydrophobicity since the change in molecular structure upon introduction of a functional group due to carbonylation is small. To account for the variation in accuracy of the predictor programs, a weight is assigned to each program based on the deviation of the prediction from our existing force field value for the amino-acids in their native state (Ghavami et al., 2014). The assigned weight (*W*_*k,i*_) for program *k* for amino acid *i* is defined as: 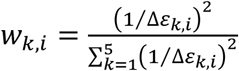 is the difference between the hydrophobicity for amino acid *i* in the 1BPA force field(Ghavami et al., 2014) and that predicted by program *k*. The results are depicted in Table S2, showing that K and R become more hydrophobic, whereas T and P become more hydrophilic, compared to their native state.

Carbonylation has additional effects. For instance, the carbonylated form of K and R, i.e. aminoadipic semialdehyde (Asa) and glutamic semialdehyde (GSA), respectively, loose their positive charge and become neutral, see Table S2 (Petrov and Zagrovic, 2011). In addition, P has a ring structure which opens up during carbonylation making the polypeptide backbone less stiff. We take this into account in our model via the bonded potential(Ghavami et al., 2013). The carbonylated form of P and R are the same (GSA)(Petrov and Zagrovic, 2011). Therefore, we assign the same hydrophobicity to them, i.e. 0.43, which is the average of the predicted hydrophobicity of 0.44 and 0.42, respectively. The relevant changes in the force field for carbonylation are summarized in Table S2.

To explore the limited effect of carbonylation on the overall distribution of the disordered phase we analyzed the changes in net hydrophobicity and charge. While the hydrophobicity for T and P is reduced, the hydrophobicity of K and R increases, resulting in only a 5% increase in net hydrophobicity for a maximally carbonylated NPC (see Table S3). Furthermore, carbonylation leads to a negatively charged NPC (−7560e) compared to a weakly positive charged wild type NPC (+512e) as all K and R become neutral. To separate the effects of charge and hydrophobicity on the structure, we carried out an additional simulation in which we consider only the change in hydrophobicity caused by carbonylation and leave the charge unaffected (termed ‘Carbonylated_HP’ in Supplementary Fig. 4c-e). The results show that the carbonylated_HP NPC is more hydrophobic than the wild type (refer to Table S3), resulting in a denser FG-Nup network with the maximum at a larger *r*-value. However, when also the charge modification is accounted for in the ‘Carbonylated’ case in Supplementary Fig. 4d the Coulombic repulsion leads to a lowering of the density, illustrating that both the change in hydrophobicity and charge affect the distribution of the disordered phase. These changes are small yet noticeable near the scaffold of the NPC, whereas the density at the center (*r* < 5 nm) is hardly affected.

### Growth of strains for oxidation assays

100 ml of BY4741 expressing Nsp1-GFP, was grown to an OD_600_ of 0.8 after which the culture was split in two portions of 50 ml; 1 portion was stressed by ROS by the addition of menadione (1 ml of 8 mg/ml in ethanol) for 1.5 hrs, while to the other only 1 ml of ethanol was added. The cells were then harvested, washed with water and stored at - 80°C until use.

### Purification of ID-GFP

*L.lactis* NZ9000 carrying a plasmid from which the yeast Heh2 ID-linker can be expressed as a GFP fusion (ID-GFP)(Meinema et al., 2011) was grown in 1 liter of GM17 medium supplemented with Chloramphenicol (5 ug/ml). When an OD600 of 0.5 was reached protein expression was induced by addition of 1 ml of the supernatant of the nisin producing *L. lactis* NZ9700. After 2 hrs the cells were harvested by centrifugation, washed once with 50 mM KPi pH7.0 and the pellet was resuspended in 5 ml of the same buffer. Drops of the suspension were frozen in liquid nitrogen and the resulting frozen droplets were pulverized in a cryomill, cooled with liquid nitrogen. 1.5 grams of the resulting powder was resuspended in 10 ml 100 mM NaPi pH7 150 mM NaCl, 10% glycerol, 0.1 mM MgCl2 5 ug/ml DnaseI, 18 mg/ml PMSF and 5 mM DTT and homogenized with a polytron (2 times 30 seconds at max speed). The suspension was cleared from non-lyzed cells by centrifugation at 20.000 rcf at 4 degrees for 20 minutes. 1 ml of Ni-sepharose slurry was pre-equilibrated in a polyprep column (BioRad) with 10 ml of ddH2O and subsequently 10 ml of 100 mM NaPi pH7 150 mM NaCl, 10% glycerol, 5 mM DTT. The cleared lysate was mixed with the equilibrated Ni-sepharose and incubated at 4 degrees under mild agitation in the polyprep column. The column was subsequently drained, washed with 10 ml 10 ml 100 mM NaPi pH7 300 mM NaCl, 10% glycerol, 15 mM imidazole, 5 mM DTT and 10 ml 10 ml 100 mM NaPi pH7 300 mM NaCl, 10% glycerol, 50 mM imidazole, 5 mM DTT. The bound ID-GFP was finally eluted from the column with 100 mM NaPi pH7 300 mM NaCl, 10% glycerol, 300 mM imidazole. The buffer was exchanged to PBS with a Zebaspin desalting column (Thermo Fisher) and the protein concentration was determined using the BCA kit (Pierce). The protein was stored overnight at 4 °C.

### Immunoprecipitation of Nsp1-GFP

Cell lysates were prepared from the cells described above in 0.5 ml lysis buffer (50 mM Kpi pH7, 250 mM NaCl, 1% Triton X100, 0.5% deoxycholate, 1 mM MgCl2, 5 mM DTT and protease inhibitors) with 0.5 mm beads in a Fastprep machine. After bead-beating the cells were incubated on ice for 15 min, and subsequently centrifuged at 20.000 rcf for 15 min to clear the lysates of beads and unbroken cells. The cell lysates were diluted with lysis buffer to 1 ml and 10 ul of GFP-nanotrap beads (Chromotek) were added. After 1.5 hrs of incubation, the beads were washed 6 times with wash buffer (50 mM Kpi pH7, 250 mM NaCl, 0.1% SDS, 0.05% Triton X100, 0.025% Deoxycholate, protease inhibitors). Bound proteins were subsequently eluted from the beads by adding 20 ul of 10% SDS to the beads and 10 min of incubation at 95 degrees. As a negative control a cell lysate from BY4741, not expressing any GFP-tagged nucleoporin, was treated as above. As positive controls BY4741 was spiked with 2 ug of purified ID-GFP, or 2 ug of ID-GFP which was first in vitro oxidized with 1 mM CuSO4 and 4 mM H_2_O_2_ for 15 min at RT.

### Western blotting and ELISA

4 ul of the eluate was separated on a 10% SDS-PAGE gel, transferred to PVDF membrane and GFP-tagged proteins were detected with anti-GFP antibodies. 10 ul of the eluate from the immunoprecipitations was used for the oxi-ELISA. First the SDS was removed from the sample using the HIPPR detergent removal kit (Pierce), by diluting the sample to 100 ul with PBS, and following the protocol as provided with the kit. The detergent free protein was subsequently diluted to 1 ml with PBS and two-fold serial dilutions were prepared in PBS. 96 well Nunc maxisorp plates (Thermo Fischer) were coated with 100 ul of the serial dilutions in duplo, and incubated O/N at 4°C. Protein amounts were determined by detection of GFP-tagged bound protein with anti-GFP antibodies and a standard ELISA protocol. In a separate ELISA plate the carbonylation state of the proteins was assessed with an Oxi-ELISA using the Oxyblott Protein oxidation detection kit (Millipore), which was essentially performed as described in Alamdari et al., 2005.

### Statistical analysis

Statistical parameters including the definitions and exact values of N, distributions and deviations are reported in the Figures and corresponding Figure legends. Significance of changes were determined with a two tailed Student’s t-test and with non-overlapping notches indicating 95% confidence that two samples are different. Unless mentioned otherwise, the experimental data coming from at least two independent cultures and microfluidic chip experiments were analysed together. In specific cases (Supplementary figure 2h and 3) the datasets deviated due to differences in filter/camera settings and are presented separately.

## Acknowledgements

We are grateful for the gift of strain MKY227 from Sanford Simon from the Rockefeller University. We thank Sara Mavrova, Jelmer de Jong, Elizabeth Carolina Riquelme Barrientos and Wouter Sipma for their help setting up the oxidation studies. We thank the Nathan Shock core at the University of Washington. We thank Sabeth Verpoorte, Michael Chang and the Veenhoff and Chang labs for their critical input into this project. This project was funded by the Netherlands Organization for Scientific Research (ECHO, NWO open to LMV) and a gift from the Ubbo Emmius Fund to the Veenhoff lab.

## Author contribution

ILR designed, performed and analysed experiments in Figure 1,2,3,4 and Supplementary Figures 2,4,5,6. AS designed, performed and analysed experiments in Supplementary Figures 3a,b and Figure 3c. AM, EG and PRO developed models in Figure 2a,b and Supplementary Figure 3cde & 8. MMC and MK designed, performed and analysed experiments in Figure 4 and Supplementary Figure 7 and designed the microfluidic chips. DPNJ, GEJ and PP were involved in generating strains and preliminary data for Figure 3. LMV and ILR wrote the manuscript with input from all authors.

---

**Supplementary Fig. 1:**
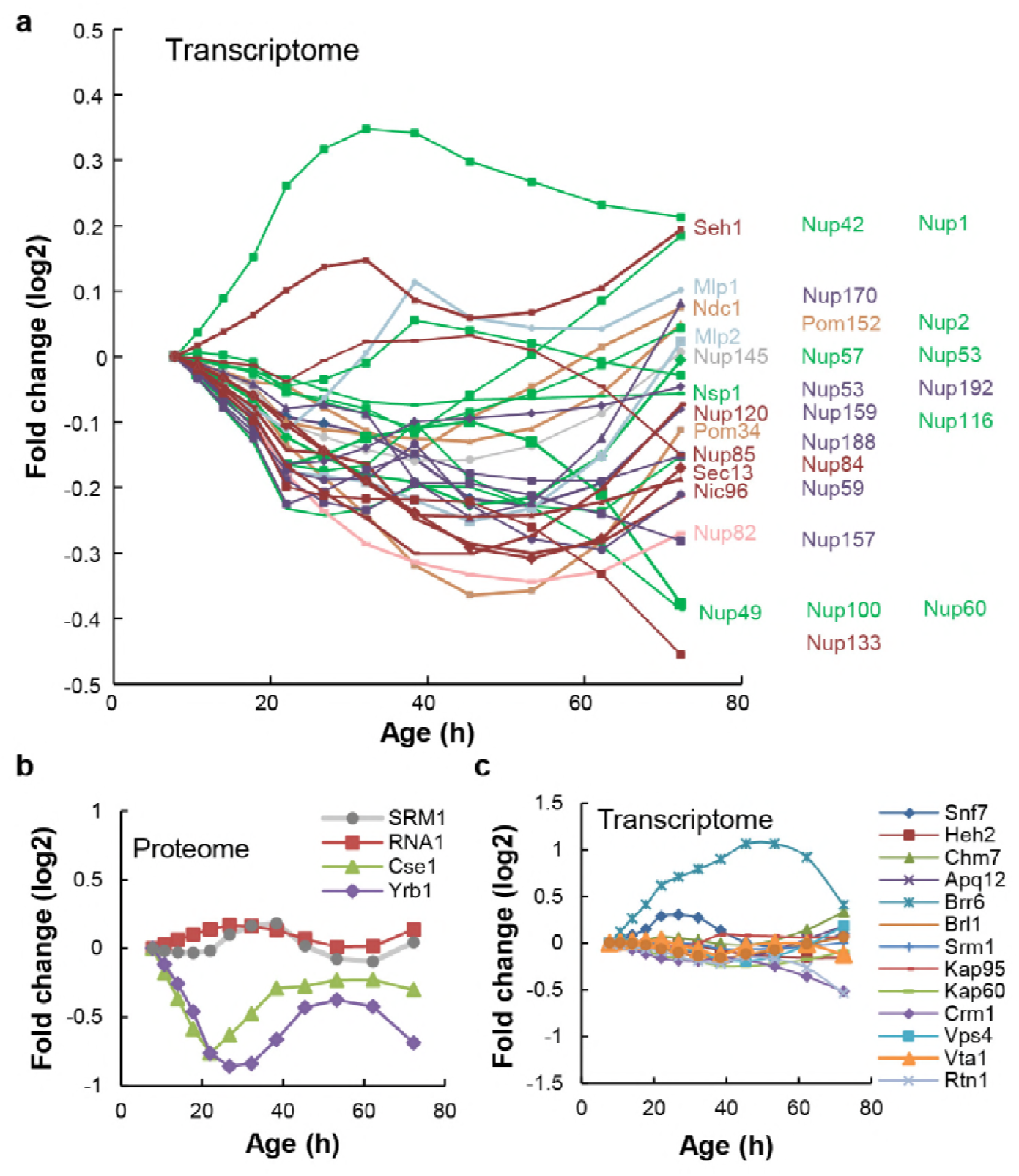
Cellular protein and mRNA abundance Nups, and NPC transport and assembly factors in replicative aging, related to Fig. 1, 2 and 3. **a,** mRNA abundance of NPC components in replicative aging. Changes in abundance are plotted as fold change. Replicative age increases in time. Transcriptome data from (Janssens et al., 2015). **b,** Protein abundance of the RanGEF Srm1, the RanGap Rna1, the RanBP1 Yrb1, and the transport receptor Cse1 as measured in whole cell extracts of yeast cells of increasing replicative age. Data from (Janssens et al., 2015). **c,** mRNA abundance of NPC assembly components and relevant NTRs (Kap95, Kap60 and Crm1) in replicative aging. Changes in abundance are plotted as fold change. Replicative age increases in time. Transcriptome data from (Janssens et al., 2015).

**Supplementary Fig. 2:**
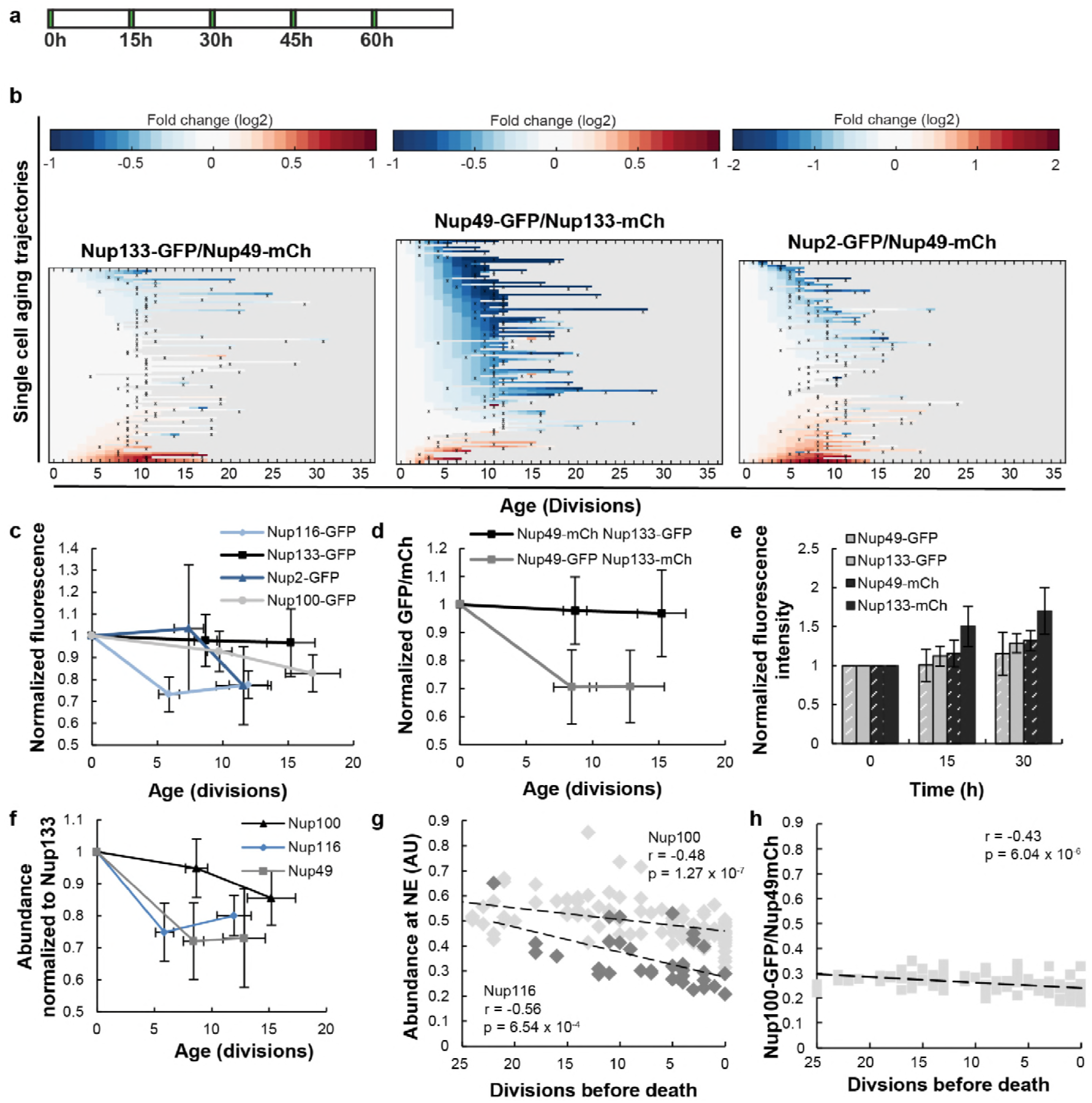
The abundance and localisation of NPC components in replicative ageing, related to Fig. 1. **a,** The experimental timeline where young cells are trapped in the microfluidic device and bright field images are taken every 20 minutes to define the cells age and fluorescent images are taken once every 15 hours to detect the protein localization and abundance. **b,** Heat map representation of the changes in the levels of the indicated GFP-and mCh-tagged Nups at the NE in each yeast cell at increasing age. Each line represents a single cell’s life history showing the change in the ratio of the fluorescence from the GFP-tagged Nup over the fluorescence of the mCh-tagged Nup and normalized to their ratio at time zero. Measurement of the fluorescence ratios are marked with “x”; in between two measurements the data was linearly interpolated. The fold changes are color coded on a log 2 scale from -1 to +1, except for Nup2 where the changes were larger and the scale runs from -2 to 2; blue colors indicate decreasing levels of the GFP-fusion relative to mCh. Number of cells in the heatmaps are Nup133-GFP/Nup49-mCh = 94, Nup49-GFP/Nup133-mCh = 108, Nup2-GFP/Nup49-mCh = 98. Data from Nup133-GFP/Nup49-mCh is repeated from Figure 1b middle panel for easy comparison. **c,** Normalized GFP/Nup49-mCh ratio representing the average from all cells shown in panel b and Fig. 1e. The indicated age is the average number of divisions after 0/15/30 h. Error bars are SD of the mean. For Nup116-GFP the change in abundance becomes significant after 15 h, with p < 0.001. For Nup2-GFP and Nup100-GFP the change in abundance is significant with p<0.005 after 30 h. The number of all measurements contributing to the means (N) at the timepoints 0 h, 15 h and 30 h were for Nup116 = 76,70 and 32; for Nup100 = 139, 137 and 86; for Nup2 = 112, 116 and 58; and for Nup133 = 102, 109 and 45, respectively. **d,** Tag-swap experiment reveals systematic changes in the fluorescence of GFP and mCh in aging: the normalized GFP/mCh ratio for Nup133-GFP/Nup49-mCh does not change significantly in aging, while the normalized GFP/mCh ratio significantly decreases (p<0.001, after 15 h) in aging for Nup49-GFP/Nup133-mCh (N = 113, 127 and 63 at timepoints 0 h, 15 h and 30 h, respectively). Error bars are SD of the mean. **e,** The average fluorescence intensities of GFP and mCh increase in time, during replicative aging experiments. The average increase in fluorescence is more pronounced in fluorophores fused to Nup133 than when fused to Nup49 showing that levels of Nup49 decrease relative to Nup133. For the strain expressing Nup49-GFP and Nup133-mCh, N = 113, 104 and 50, and for the strains expressing Nup133-GFP and Nup49-mCh, N = 102, 85 and 27 after 0 h, 15 h and 30 h respectively. Error bars are SD of the mean. **f,** Abundance of Nup100-GFP, Nup116-GFP and Nup49-GFP, relative to Nup133-GFP. **g,** The abundance of Nup116-GFP (grey) and Nup100 (black) at the NE as a function of remaining lifespan. The dotted lines indicate the best linear fit. Total number of cells analyzed are Nup116 = 15 and Nup100 = 35 and the total number of measurements are Nup116 = 34 and Nup100 = 108. **h,** Additional independent replicate (coming from a different microscope) for Nup100-GFP/Nup49-mCh abundance correlation to lifespan. The cells in h and g were imaged with different filter settings explaining the different ratios. Number of cells analyzed are N = 62 and number of measurements are N = 101.

**Supplementary Fig. 3:**
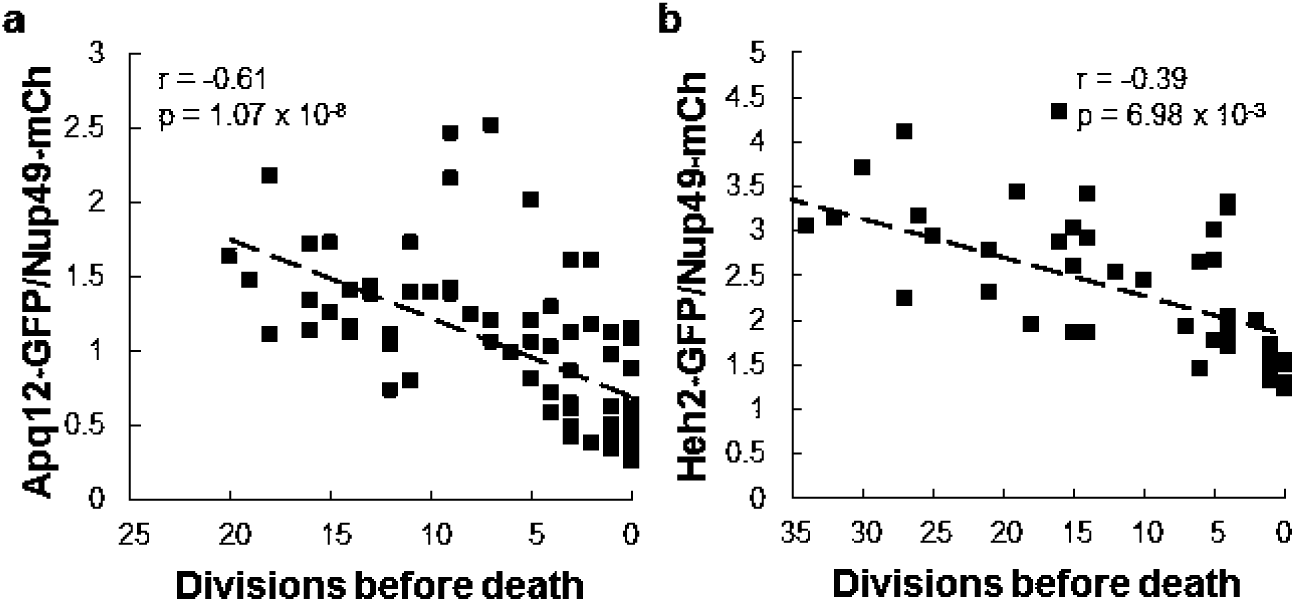
Heh2-GFP and Apq12-GFP abundance at the NE as a function of remaining lifespan related to Fig. 2. Additional independent replicate (coming from a different microscope) for Apq12-GFP and Heh2-GFP abundance at the NE, relative to Nup49-mCh, as a function of remaining lifespan. The dotted lines indicate best linear fit; Pearson correlations are indicated. Number of cells analyzed are Apq12 = 34, Heh2 = 14 and number of measuring points analyzed are Apq12 = 74 and Heh2 = 46.

**Supplementary Fig. 4:**
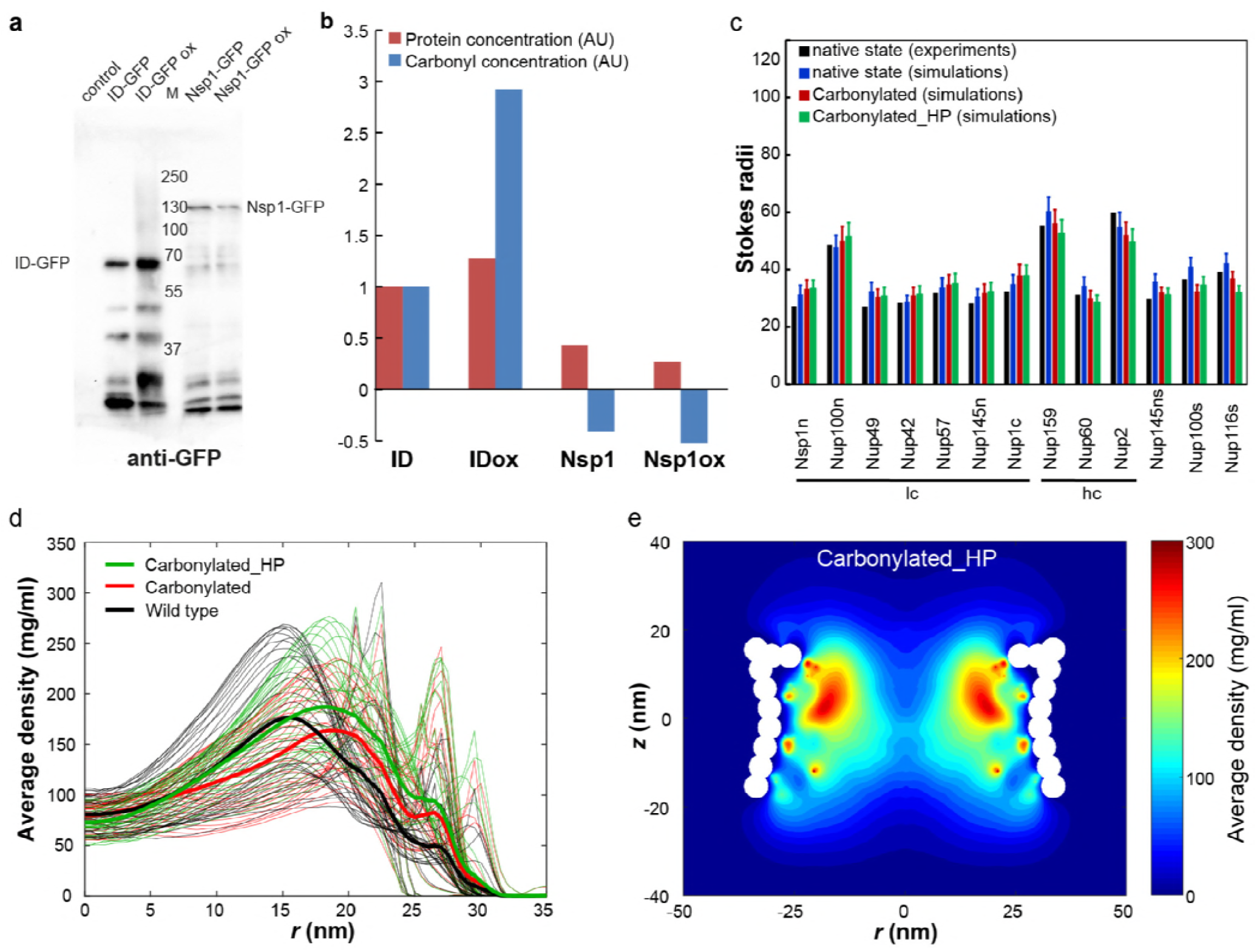
In vitro oxidation and models of NPCs with oxidative damage, related to Fig. 2. **a,** Anti-GFP Western blot of Nsp1-GFP immunoprecititated from extracts of exponentially growing BY4741 cell expressing Nsp1-GFP from the native promotor and treated without or with 1 mM menadione for 90 minutes to induce high ROS levels (Nsp1-GFP, Nsp1-GFP ox). As positive control in vitro oxidized purified ID protein (namely the ID linker of Heh2 fused to GFP, ID-GFPox) was added to the BY4741 cell extracts before immunoprecipitation. BY4741 cell extract with and without additions of non-oxidized ID-GFP serve as negative controls (ID-GFP and control). **b,** Carbonyl detection with ELISA. The immunoprecipitated samples were cleaned from detergents, and serial dilutions were bound to Nunc maxisorp ELISA plates and an ELISA with GFP antibodies as well as an oxi-ELISA, essentially as described by (Alamdari et al., 2005), were performed. The read outs of both ELISAs represent the amount of protein (anti-GFP, red bars) or carbonyls (oxi-ELISA with anti-DNP, blue bars) on ID-GFP and Nsp1; carbonyl levels on Nsp1-GFP are below the detection level even under these strongly oxidizing conditions. **c,** The Stokes radii for FG-Nups and FG-Nup segments for the native and carbonylated state (in Angstrom). The black bar represents the experimental (native) Stokes radii from (Yamada et al., 2010), the blue bar represents the prediction for these native FG Nups (Ghavami et al., 2014), the prediction for the carbonylated FG Nups is plotted in red and the results for the carbonylated_HP (see Methods for details) variant is shown in green. The error bar for the simulations represents the standard deviation in time. **d,** Time averaged radial density plot for a carbonylated_HP NPC compared with the wild type and carbonylated NPCs at different positions along the *z*-axis separated by 1 nm in the *z*-range of -15.4 to 15.4 nm. In the carbonylated_HP NPC only the effect of carbonylation on the hydrophobicity is accounted for. The average over the different *z*-values is plotted as thick lines for all three cases. **e,** Two-dimensional (*rz*) density map for the carbonylated_HP NPC.

**Supplementary Fig. 5:**
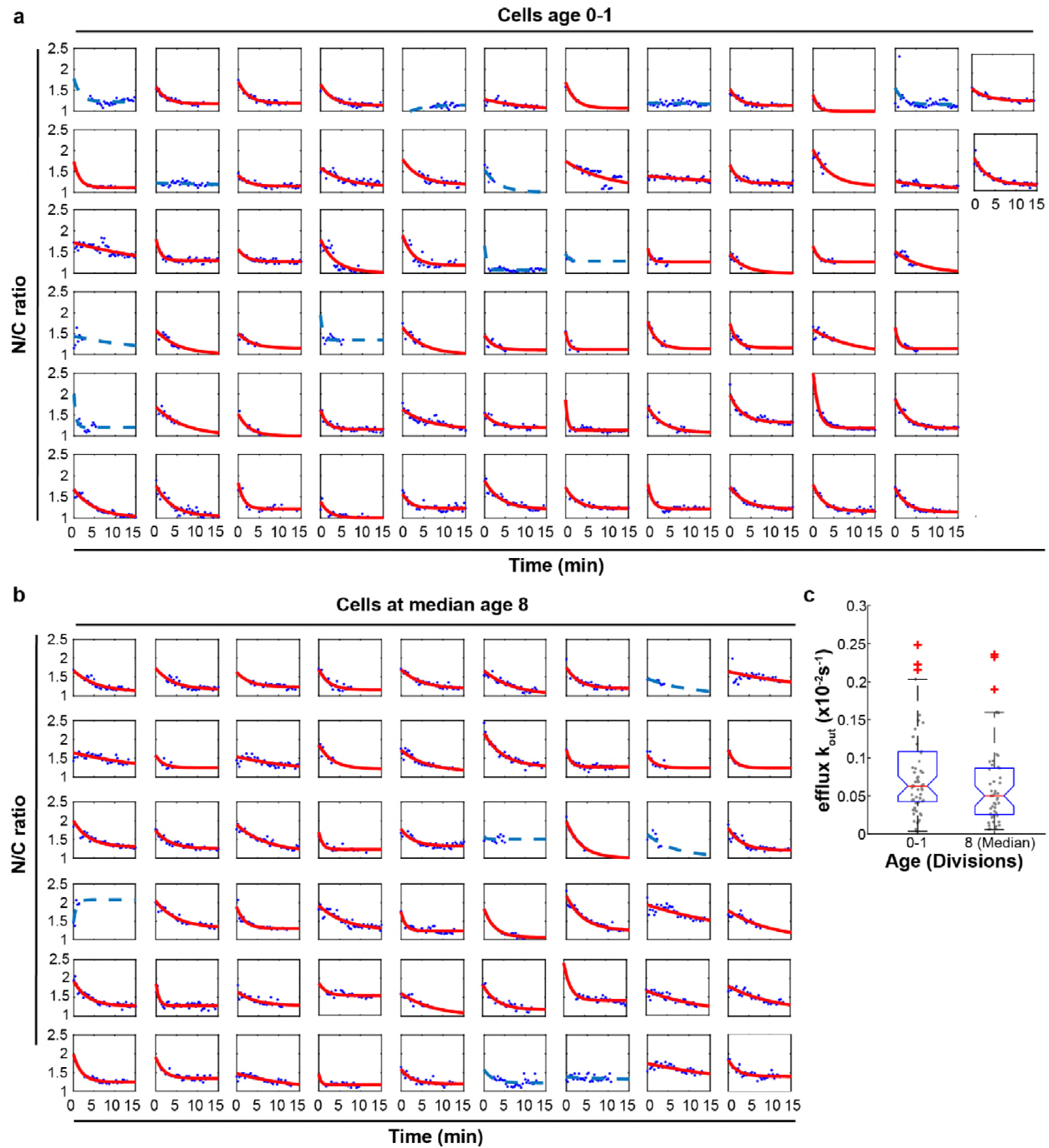
Efflux rate constants and NTR localisation in ageing, related to Fig. 3. **a,b,** Singe cell measurements of the kinetics of loss of nuclear accumulation of GFP-NLS from young cell and cell with median replicative age 8. Time zero is the time the medium with Na-azide and 2-Deoxy-D-glucose and ponceau red for visibility reached the cells, trapped in the device. The measurements are fitted to an exponential decay function and yield the efflux rate constant (k_out_). Only cells with p < 0.05 and R^2^>0.2 (plotted in red) are represented in panel c; poor fits (blue lines) are excluded from the analysis. **c,** Efflux rate constant of cells age 0 and cells age 8. The average K_out_ of old cells is lower than for young cells, but changes are not significant. Number of cells included in the analysis are Age 0-1 = 57 and 8 (Median) = 48.

**Supplementary Fig. 6:**
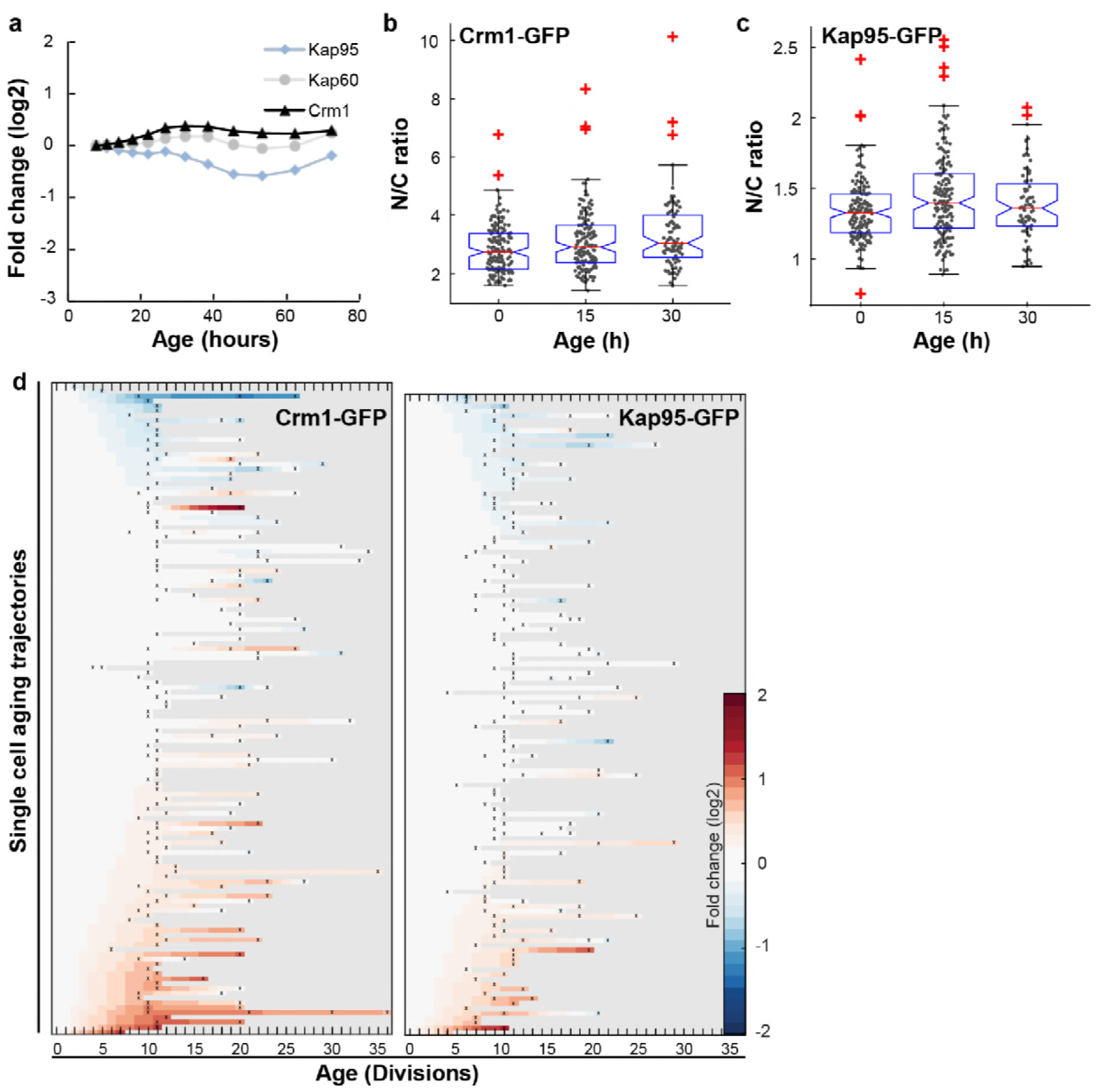
The abundance and localization of transport factors does not change in aging, related to Fig. 3. **a,** Protein abundance of Crm1, Kap95 and Kap60 as measured in whole cell extracts of yeast cells of increasing replicative age. Data from (Janssens et al., 2015). **b,c,** Localization of Crm1 b, and Kap95 c, during replicative aging to the nucleus relative to the cytosol (N/C ratio). The line indicates the median, and the bottom and top edges of the box indicate the 25th and 75th percentiles, respectively. The whiskers extend to the most extreme data points not considered outliers, and the outliers are plotted individually. Non overlapping notches indicate that the samples are different with 95% confidence. The overall changes were thus not significant, although we note that based on a two-tailed Student’s T-test the N/C ratio for Kap95 is significantly increased after 15 h (p = 8.7 x 10^-4^). No significant correlation was found with age (Crm1: r = 0.15, p = 0.09 and Kap95: r = 0.07, p = 0.39), or lifespan (Crm1: r = 0.04, p = 0.63 and Kap95: r = 0.11, p = 0.16). Number of cells analyzed at 0 h, 15 h, and 30 h were for Kap95 = 155, 165, 72 and for Crm1 = 156, 138, 87. **d,** Heatmap representation of changes in N/C ratio of Crm1-GFP (N = 134) and Kap95-GFP (N = 132).

**Supplementary Fig. 7:**
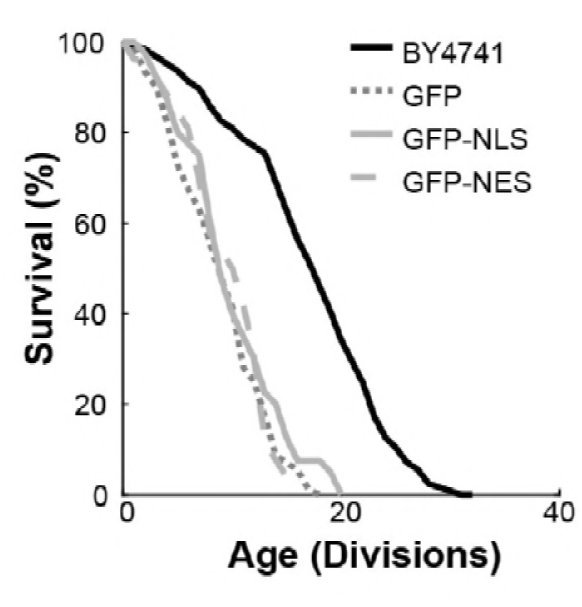
Replicative lifespan curves, related to Fig. 3. Replicative lifespan curves of strains expressing reporter proteins, in comparison to BY4741. The expression of GFP alone did not result in any observable growth defect in young cells, but did impact the lifespan of the yeast cells. This impact on lifespan is likely related to a general stress resulting from the additional protein synthesis and is unlikely to be related to nuclear transport. To enable comparison of the three reporter proteins, GFP, GFP-NLS and GFP-NES, in aging, we tuned their expression such that the impact on lifespan was similar for all three. Total number of cells analysed per strain were GFP = 89, GFP-NLS = 96, GFP-NES = 75, BY4741 = 126.

**Supplementary Fig. 8:**
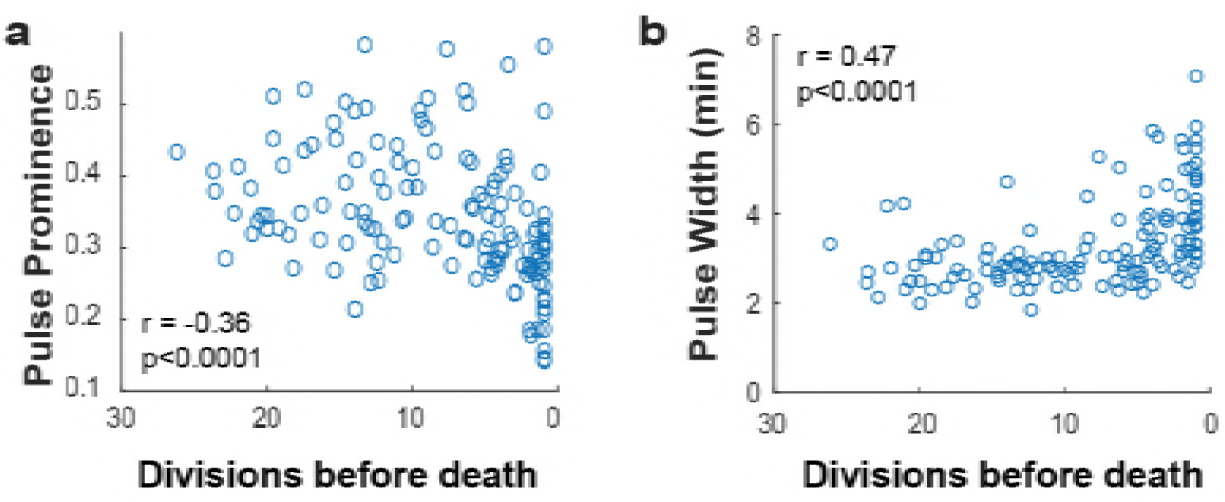
Msn2 pulse prominence and width correlate to remaining lifespan, related to Fig. 4. **a,** The Msn2 pulse prominence is positively correlated with remaining lifespan. **b,** The remaining lifespan is negatively correlated with increasing Msn2 pulse width. N value cells = 48, N value for scatter plot = 138

**Supplementary Fig. 9:**
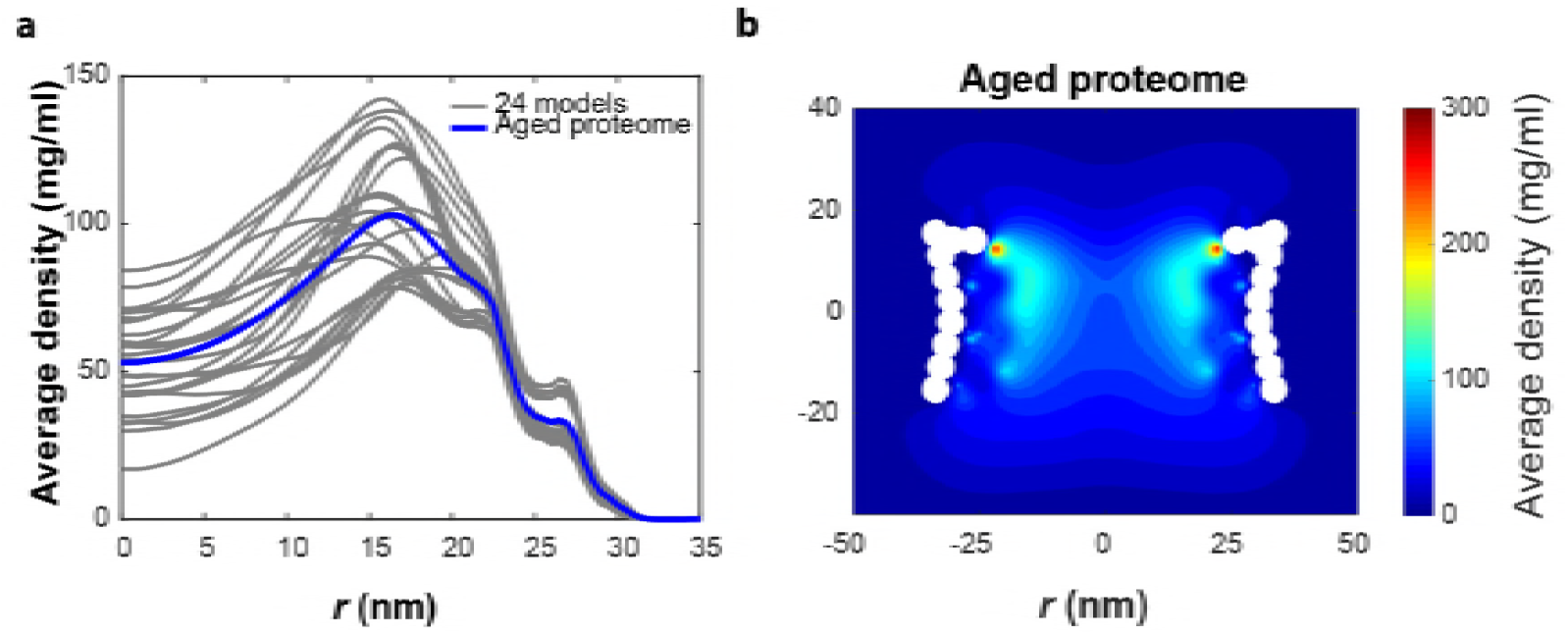
Models of NPCs with altered stoichiometry. **a,** The time averaged radial density distribution of the 24 constructed models (grey) (see Table S1) based on the FG-Nup abundance data from Fig. 1c with the average of the 24 models plotted in blue denoted as ‘Aged proteome’. Each curve in grey represents the radial density averaged over the NPC height i.e. |*z|* < 15.4 nm for one of the 24 different models. **b,** Time-averaged *r-z* density of FG-Nups the aged proteome NPC.

**Table S1:** Details of the FG-Nup stoichiometry for the 24 constructed models to represent the aged NPC. 0 and 1 represent absence and presence, respectively, of the FG-Nup in 8-fold symmetry.

|  | NSP1 | NSP1 | NSP1 | NSP1 | Nup1 | Nup42 | Nup49 | Nup49 | Nup57 | Nup57 | Nup60 | Nup100 | Nup145 | Nup145 | Nup159 | Nup116 |
| --- | --- | --- | --- | --- | --- | --- | --- | --- | --- | --- | --- | --- | --- | --- | --- | --- |
| model 1 | 0 | 0 | 1 | 1 | 1 | 1 | 1 | 1 | 1 | 1 | 1 | 1 | 1 | 1 | 1 | 0 |
| model 2 | 0 | 0 | 1 | 1 | 1 | 1 | 1 | 1 | 1 | 1 | 1 | 0 | 1 | 1 | 1 | 0 |
| model 3 | 0 | 0 | 1 | 1 | 1 | 1 | 1 | 1 | 1 | 1 | 0 | 1 | 1 | 1 | 1 | 0 |
| model 4 | 0 | 0 | 1 | 1 | 1 | 1 | 1 | 1 | 1 | 1 | 0 | 0 | 1 | 1 | 1 | 0 |
| model 5 | 0 | 0 | 1 | 1 | 0 | 1 | 1 | 1 | 1 | 1 | 1 | 1 | 1 | 1 | 1 | 0 |
| model 6 | 0 | 0 | 1 | 1 | 0 | 1 | 1 | 1 | 1 | 1 | 1 | 0 | 1 | 1 | 1 | 0 |
| model 7 | 0 | 0 | 1 | 1 | 0 | 1 | 1 | 1 | 1 | 1 | 0 | 1 | 1 | 1 | 1 | 0 |
| model 8 | 0 | 0 | 1 | 1 | 0 | 1 | 1 | 1 | 1 | 1 | 0 | 0 | 1 | 1 | 1 | 0 |
| model 9 | 0 | 0 | 1 | 0 | 1 | 1 | 0 | 1 | 0 | 1 | 1 | 1 | 1 | 1 | 1 | 0 |
| model 10 | 0 | 0 | 1 | 0 | 1 | 1 | 0 | 1 | 0 | 1 | 1 | 0 | 1 | 1 | 1 | 0 |
| model 11 | 0 | 0 | 1 | 0 | 1 | 1 | 0 | 1 | 0 | 1 | 0 | 1 | 1 | 1 | 1 | 0 |
| model 12 | 0 | 0 | 1 | 0 | 1 | 1 | 0 | 1 | 0 | 1 | 0 | 0 | 1 | 1 | 1 | 0 |
| model 13 | 0 | 0 | 1 | 0 | 0 | 1 | 0 | 1 | 0 | 1 | 1 | 1 | 1 | 1 | 1 | 0 |
| model 14 | 0 | 0 | 1 | 0 | 0 | 1 | 0 | 1 | 0 | 1 | 1 | 0 | 1 | 1 | 1 | 0 |
| model 15 | 0 | 0 | 1 | 0 | 0 | 1 | 0 | 1 | 0 | 1 | 0 | 1 | 1 | 1 | 1 | 0 |
| model 16 | 0 | 0 | 1 | 0 | 0 | 1 | 0 | 1 | 0 | 1 | 0 | 0 | 1 | 1 | 1 | 0 |
| model 17 | 0 | 0 | 0 | 1 | 1 | 1 | 1 | 0 | 1 | 0 | 1 | 1 | 1 | 1 | 1 | 0 |
| model 18 | 0 | 0 | 0 | 1 | 1 | 1 | 1 | 0 | 1 | 0 | 1 | 0 | 1 | 1 | 1 | 0 |
| model 19 | 0 | 0 | 0 | 1 | 1 | 1 | 1 | 0 | 1 | 0 | 0 | 1 | 1 | 1 | 1 | 0 |
| model 20 | 0 | 0 | 0 | 1 | 1 | 1 | 1 | 0 | 1 | 0 | 0 | 0 | 1 | 1 | 1 | 0 |
| model 21 | 0 | 0 | 0 | 1 | 0 | 1 | 1 | 0 | 1 | 0 | 1 | 1 | 1 | 1 | 1 | 0 |
| model 22 | 0 | 0 | 0 | 1 | 0 | 1 | 1 | 0 | 1 | 0 | 1 | 0 | 1 | 1 | 1 | 0 |
| model 23 | 0 | 0 | 0 | 1 | 0 | 1 | 1 | 0 | 1 | 0 | 0 | 1 | 1 | 1 | 1 | 0 |
| model 24 | 0 | 0 | 0 | 1 | 0 | 1 | 1 | 0 | 1 | 0 | 0 | 0 | 1 | 1 | 1 | 0 |

**Table S2:** Force field parameters for carbonylated amino acids. Here *ε*_Ghavami_ and *ε*_native_ represent the hydrophobicity of amino acids in their native condition according to (Ghavami et al., 2014) and the weighted average scheme, respectively. *ε*_carbonylated_ denotes the hydrophobicity derived from the weighted average scheme and *q*_native to carbonylated_ stands for the charge modification from the native to the carbonylated state.

| AA | $\epsilon_{\text{Ghavami}}$ | $\epsilon_{\text{native}}$ | $\epsilon_{\text{carbonylated}}$ | $q_{\text{native to carbonylated}}$ |
| --- | --- | --- | --- | --- |
| T | 0.51 | 0.52 | 0.34 | 0 → 0 |
| K | 0.00 | 0.00 | 0.59 | 1 → 0 |
| P | 0.65 | 0.63 | 0.43 | 0 → 0 |
| R | 0.00 | 0.07 | 0.43 | 1 → 0 |

**Table S3:** Physical properties of the wild type, carbonylated and carbonylated_HP NPCs. In the carbonylated_HP NPC only the effect of carbonylation on the hydrophobicity is accounted for. For the net hydrophobicity we added the hydrophobicity values of all the residues inside the NPC.

| Force field | +ve Charged AA | -ve Charged AA | Net charge | Net hydrophobicity |
| --- | --- | --- | --- | --- |
| Wild Type | 8072 | 7560 | +512 | 43373.7 |
| Carbonylated | 0 | 7560 | -7560 | 45549.7 |
| Carbonylated_HP | 8072 | 7560 | +512 | 45549.7 |

## References

Alamdari, D.H., Kostidou, E., Paletas, K., Sarigianni, M., Konstas, A.G.P., Karapiperidou, A., and Koliakos, G. (2005). High sensitivity enzyme-linked immunosorbent assay (ELISA) method for measuring protein carbonyl in samples with low amounts of protein. Free Radic. Biol. Med. 39, 1362–1367.

Alber, F., Dokudovskaya, S., Veenhoff, L.M., Zhang, W., Kipper, J., Devos, D., Suprapto, A., Karni-Schmidt, O., Williams, R., Chait, B.T., et al. (2007). The molecular architecture of the nuclear pore complex. Nature 450, 695–701.

Bakker, E., Swain, P.S., and Crane, M.M. (2018). Morphologically constrained and data informed cell segmentation of budding yeast. Bioinformatics 34, 88–96.

Cai, L., Dalal, C.K., and Elowitz, M.B. (2008). Frequency-modulated nuclear localization bursts coordinate gene regulation. Nature 455, 485–490.

Chadrin, A., Hess, B., San Roman, M., Gatti, X., Lombard, B., Loew, D., Barral, Y., Palancade, B., and Doye, V. (2010). Pom33, a novel transmembrane nucleoporin required for proper nuclear pore complex distribution. J. Cell Biol. 189, 795–811.

Colombi, P., Webster, B.M., Fröhlich, F., and Patrick Lusk, C. (2013). The transmission of nuclear pore complexes to daughter cells requires a cytoplasmic pool of Nsp1. J. Cell Biol.

Crane, M.M., Clark, I.B.N., Bakker, E., Smith, S., and Swain, P.S. (2014). A microfluidic system for studying ageing and dynamic single-cell responses in budding yeast. PLoS One 9.

Cutler, A.A., Dammer, E.B., Doung, D.M., Seyfried, N.T., Corbett, A.H., and Pavlath, G.K. (2017). Biochemical isolation of myonuclei employed to define changes to the myonuclear proteome that occur with aging. Aging Cell 16, 738–749.

D’Angelo, M.A., Raices, M., Panowski, S.H., and Hetzer, M.W. (2009). Age-Dependent Deterioration of Nuclear Pore Complexes Causes a Loss of Nuclear Integrity in Postmitotic Cells. Cell 136, 284–295.

Dawson, T.R., Lazarus, M.D., Hetzer, M.W., and Wente, S.R. (2009). ER membrane-bending proteins are necessary for de novo nuclear pore formation. J. Cell Biol. 184, 659–675.

Denoth-Lippuner, A., Krzyzanowski, M.K., Stober, C., and Barral, Y. (2014). Role of SAGA in the asymmetric segregation of DNA circles during yeast ageing. Elife.

Denoth Lippuner, A., Julou, T., and Barral, Y. (2014). Budding yeast as a model organism to study the effects of age. FEMS Microbiol. Rev. 38, 300–325.

Dilworth, D.J., Suprapto, A., Padovan, J.C., Chait, B.T., Wozniak, R.W., Rout, M.P., and Aitchison, J.D. (2001). Nup2p dynamically associates with the distal regions of the yeast nuclear pore complex. J. Cell Biol. 153, 1465–1478.

Fehrmann, S., Paoletti, C., Goulev, Y., Ungureanu, A., Aguilaniu, H., and Charvin, G. (2013). Aging yeast cells undergo a sharp entry into senescence unrelated to the loss of mitochondrial membrane potential. Cell Rep.

Fichtman, B., and Harel, A. (2014). Stress and aging at the nuclear gateway. Mech. Ageing Dev. 135, 24–32.

Fiserova, J., and Goldberg, M.W. (2010). Nucleocytoplasmic transport in yeast: a few roles for many actors. Biochem. Soc. Trans. 38, 273–277.

Fujita, T., Iwasa, J., and Hansch, C. (1964). A New Substituent Constant, π, Derived from Partition Coefficients. J. Am. Chem. Soc. 86, 5175–5180.

Ghavami, A., van der Giessen, E., and Onck, P.R. (2013). Coarse-Grained Potentials for Local Interactions in Unfolded Proteins. J. Chem. Theory Comput. 9, 432–440.

Ghavami, A., Veenhoff, L.M., Van Der Giessen, E., and Onck, P.R. (2014). Probing the disordered domain of the nuclear pore complex through coarse-grained molecular dynamics simulations. Biophys. J.

Granados, A.A., Crane, M.M., Montano-Gutierrez, L.F., Tanaka, R.J., Voliotis, M., and Swain, P.S. (2017). Distributing tasks via multiple input pathways increases cellular survival in stress. Elife 6.

Hao, N., and O’Shea, E.K. (2011). Signal-dependent dynamics of transcription factor translocation controls gene expression. Nat. Struct. Mol. Biol. 19, 31–39.

Hao, N., Budnik, B.A., Gunawardena, J., and O’Shea, E.K. (2013). Tunable signal processing through modular control of transcription factor translocation. Science 339, 460–464.

Huh, W.-K., Falvo, J. V, Gerke, L.C., Carroll, A.S., Howson, R.W., Weissman, J.S., and O’shea, E.K. Global analysis of protein localization in budding yeast.

Hurt, E., and Beck, M. (2015). Towards understanding nuclear pore complex architecture and dynamics in the age of integrative structural analysis. Curr. Opin. Cell Biol.

Janke, C., Magiera, M.M., Rathfelder, N., Taxis, C., Reber, S., Maekawa, H., Moreno-Borchart, A., Doenges, G., Schwob, E., Schiebel, E., et al. (2004). A versatile toolbox for PCR-based tagging of yeast genes: New fluorescent proteins, more markers and promoter substitution cassettes. Yeast.

Janssens, G., and Veenhoff, L. (2016a). Evidence for the hallmarks of human aging in replicatively aging yeast. Microb. Cell 3, 263–274.

Janssens, G.E., and Veenhoff, L.M. (2016b). The Natural Variation in Lifespans of Single Yeast Cells Is Related to Variation in Cell Size, Ribosomal Protein, and Division Time. PLoS One 11, e0167394.

Janssens, G.E., Meinema, A.C., Gonz??lez, J., Wolters, J.C., Schmidt, A., Guryev, V., Bischoff, R., Wit, E.C., Veenhoff, L.M., and Heinemann, M. (2015). Protein biogenesis machinery is a driver of replicative aging in yeast. Elife 4.

Jo, M.C., Liu, W., Gu, L., Dang, W., and Qin, L. High-throughput analysis of yeast replicative aging using a microfluidic system.

Khmelinskii, A., Keller, P.J., Bartosik, A., Meurer, M., Barry, J.D., Mardin, B.R., Kaufmann, A., Trautmann, S., Wachsmuth, M., Pereira, G., et al. (2012). Tandem fluorescent protein timers for in vivo analysis of protein dynamics.

Kim, S.J., Fernandez-Martinez, J., Nudelman, I., Shi, Y., Zhang, W., Raveh, B., Herricks, T., Slaughter, B.D., Hogan, J.A., Upla, P., et al. (2018). Integrative structure and functional anatomy of a nuclear pore complex. Nature 555, 475–482.

Lee, S.S., Vizcarra, I.A., Huberts, D.H.E.W., Lee, L.P., and Heinemann, M. Whole lifespan microscopic observation of budding yeast aging through a microfluidic dissection platform.

Leo, A.J. (1993). Calculating log Poct from structures. Chem. Rev. 93, 1281–1306.

Li, H.Y., Wirtz, D., and Zheng, Y. (2003). A mechanism of coupling RCC1 mobility to RanGTP production on the chromatin in vivo. J. Cell Biol. 160, 635–644.

Lone, M.A., Atkinson, A.E., Hodge, C.A., Cottier, S., Martínez-Montañés, F., Maithel, S., Mène-Saffrané, L., Cole, C.N., and Schneiter, R. (2015). Yeast integral membrane proteins Apq12, Brl1, and Brr6 form a complex important for regulation of membrane homeostasis and nuclear pore complex biogenesis. Eukaryot. Cell 14, 1217–1227.

Longo, V.D., Shadel, G.S., Kaeberlein, M., and Kennedy, B. (2012). Replicative and chronological aging in saccharomyces cerevisiae. Cell Metab. 16, 18–31.

Lord, C.L., Timney, B.L., Rout, M.P., and Wente, S.R. (2015). Altering nuclear pore complex function impacts longevity and mitochondrial function in S. cerevisiae. J. Cell Biol. 208, 729–744.

Maisonneuve, E., Ducret, A., Khoueiry, P., Lignon, S., Longhi, S., Talla, E., and Dukan, S. (2009). Rules governing selective protein carbonylation. PLoS One 4, e7269.

Mattheyses, A.L., Kampmann, M., Atkinson, C.E., and Simon, S.M. (2010). Fluorescence anisotropy reveals order and disorder of protein domains in the nuclear pore complex. Biophys. J. 99, 1706–1717.

Meinema, A.C., Laba, J.K., Hapsari, R.A., Otten, R., Mulder, F.A.A., Kralt, A., van den Bogaart, G., Lusk, C.P., Poolman, B., and Veenhoff, L.M. (2011). Long Unfolded Linkers Facilitate Membrane Protein Import Through the Nuclear Pore Complex. Science (80-.). 333, 90–93.

Meinema, A.C., Poolman, B., and Veenhoff, L.M. (2013). Quantitative Analysis of Membrane Protein Transport Across the Nuclear Pore Complex. Traffic 14, 487–501.

Meylan, W.M., and Howard, P.H. (1995). Atom/fragment contribution method for estimating octanol-water partition coefficients. J. Pharm. Sci. 84, 83–92.

Nemergut, M.E., Mizzen, C.A., Stukenberg, T., Allis, C.D., and Macara, I.G. (2001). Chromatin docking and exchange activity enhancement of RCC1 by histones H2A and H2B. Science 292, 1540–1543.

Niño, C.A., Guet, D., Gay, A., Brutus, S., Jourquin, F., Mendiratta, S., Salamero, J., Géli, V., and Dargemont, C. (2016). Posttranslational marks control architectural and functional plasticity of the nuclear pore complex basket. J. Cell Biol. 212, 167–180.

Nyström, T., and Liu, B. (2014). The mystery of aging and rejuvenation-a budding topic. Curr. Opin. Microbiol.

Onischenko, E., Tang, J.H., Andersen, K.R., Knockenhauer, K.E., Vallotton, P., Derrer, C.P., Kralt, A., Mugler, C.F., Chan, L.Y., Schwartz, T.U., et al. (2017). Natively Unfolded FG Repeats Stabilize the Structure of the Nuclear Pore Complex. Cell 171, 904–917.e19.

Ori, A., Toyama, B.H., Harris, M.S., Bock, T., Iskar, M., Bork, P., Ingolia, N.T., Hetzer, M.W., and Beck, M. (2015). Integrated Transcriptome and Proteome Analyses Reveal Organ-Specific Proteome Deterioration in Old Rats. Cell Syst. 1, 224–237.

Otsuka, S., and Ellenberg, J. (2018). Mechanisms of nuclear pore complex assembly - two different ways of building one molecular machine. FEBS Lett. 592, 475–488.

Petrov, D., and Zagrovic, B. (2011). Microscopic analysis of protein oxidative damage: effect of carbonylation on structure, dynamics, and aggregability of villin headpiece. J. Am. Chem. Soc. 133, 7016–7024.

Popken, Ghavami, A., Onck, P.R., Poolman, B., and Veenhoff, L.M. (2015). Size-Dependent Leak of Soluble and Membrane Proteins Through the Yeast Nuclear Pore Complex. Mol. Biol. Cell 26, 1386–1394.

Rabut, G., Doye, V., and Ellenberg, J. (2004). Mapping the dynamic organization of the nuclear pore complex inside single living cells. Nat. Cell Biol. 6, 1114–1121.

Riddick, G., and Macara, I.G. (2005). A systems analysis of importin-{alpha}-{beta} mediated nuclear protein import. J. Cell Biol. 168, 1027–1038.

Savas, J.N., Toyama, B.H., Xu, T., Yates, J.R., and Hetzer, M.W. (2012). Extremely long-lived nuclear pore proteins in the rat brain. Science (80-.). 335, 942.

Scarcelli, J.J., Hodge, C.A., and Cole, C.N. (2007). The yeast integral membrane protein Apq12 potentially links membrane dynamics to assembly of nuclear pore complexes. J. Cell Biol. 178, 799–812.

Schindelin, J., Arganda-Carreras, I., Frise, E., Kaynig, V., Longair, M., Pietzsch, T., Preibisch, S., Rueden, C., Saalfeld, S., Schmid, B., et al. (2012). Fiji: an open-source platform for biological-image analysis. Nat. Methods 9, 676–682.

Shulga, N., Roberts, P., Gu, Z., Spitz, L., Tabb, M.M., Nomura, M., and Goldfarb, D.S. (1996). In vivo nuclear transport kinetics in Saccharomyces cerevisiae: a role for heat shock protein 70 during targeting and translocation. J. Cell Biol. 135, 329–339.

Smith, A.E., Slepchenko, B.M., Schaff, J.C., Loew, L.M., and Macara, I.G. (2002). Systems Analysis of Ran Transport. Science (80-.). 295, 488–491.

Stadtman, E.R., and Levine, R.L. (2003). Free radical-mediated oxidation of free amino acids and amino acid residues in proteins. Amino Acids 25, 207–218.

Strawn, L.A., Shen, T., Shulga, N., Goldfarb, D.S., and Wente, S.R. (2004). Minimal nuclear pore complexes define FG repeat domains essential for transport. Nat. Cell Biol. 6, 197–206.

Teimer, R., Kosinski, J., von Appen, A., Beck, M., and Hurt, E. (2017). A short linear motif in scaffold Nup145C connects Y-complex with pre-assembled outer ring Nup82 complex. Nat. Commun. 8, 1107.

Tetko, I. V., Gasteiger, J., Todeschini, R., Mauri, A., Livingstone, D., Ertl, P., Palyulin, V.A., Radchenko, E. V., Zefirov, N.S., Makarenko, A.S., et al. (2005). Virtual Computational Chemistry Laboratory – Design and Description. J. Comput. Aided. Mol. Des. 19, 453–463.

Thaller, D.J., and Patrick Lusk, C. (2018). Fantastic nuclear envelope herniations and where to find them. Biochem. Soc. Trans. BST20170442.

Thayer, N.H., Leverich, C.K., Fitzgibbon, M.P., Nelson, Z.W., Henderson, K.A., Gafken, P.R., Hsu, J.J., Gottschling, D.E., Marcotte, E.M., and Denic, V. Identification of long-lived proteins retained in cells undergoing repeated asymmetric divisions.

Timney, B.L., Raveh, B., Mironska, R., Trivedi, J.M., Kim, S.J., Russel, D., Wente, S.R., Sali, A., and Rout, M.P. (2016). Simple rules for passive diffusion through the nuclear pore complex. J. Cell Biol. 215, 57–76.

Tkach, J.M., Yimit, A., Lee, A.Y., Riffle, M., Costanzo, M., Jaschob, D., Hendry, J.A., Ou, J., Moffat, J., Boone, C., et al. (2012). Dissecting DNA damage response pathways by analysing protein localization and abundance changes during DNA replication stress. Nat. Cell Biol. 14, 966–976.

Toyama, B.H., Savas, J.N., Park, S.K., Harris, M.S., Ingolia, N.T., Yates, J.R., and Hetzer, M.W. (2013). Identification of long-lived proteins reveals exceptional stability of essential cellular structures. Cell.

Viswanadhan, V.N., Ghose, A.K., Revankar, G.R., and Robins, R.K. (1989). Atomic physicochemical parameters for three dimensional structure directed quantitative structure-activity relationships. 4. Additional parameters for hydrophobic and dispersive interactions and their application for an automated superposition of certain naturally occurring nucleoside antibiotics. J. Chem. Inf. Model. 29, 163–172.

Webster, B.M., Colombi, P., Jäger, J., and Patrick Lusk, C. (2014). Surveillance of nuclear pore complex assembly by ESCRT-III/Vps4. Cell.

Webster, B.M., Thaller, D.J., Jäger, J., Ochmann, S.E., Borah, S., Lusk, C.P., Adell, M., Vogel, G., Pakdel, M., Müller, M., et al. (2016). Chm7 and Heh1 collaborate to link nuclear pore complex quality control with nuclear envelope sealing. EMBO J. 205, 695–701.

Wente, S.R., and Blobel, G. A Temperature-sensitive NUPll6 Null Mutant Forms a Nuclear Envelope Seal over the Yeast Nuclear Pore Complex Thereby Blocking Nucleocytoplasmic Traffic.

Yamada, J., Phillips, J.L., Patel, S., Goldfien, G., Calestagne-Morelli, A., Huang, H., Reza, R., Acheson, J., Krishnan, V. V, Newsam, S., et al. (2010). A bimodal distribution of two distinct categories of intrinsically disordered structures with separate functions in FG nucleoporins. Mol. Cell. Proteomics 9, 2205–2224.

Zhang, W., Neuner, A., Rüthnick, D., Sachsenheimer, T., Lüchtenborg, C., Brügger, B., and Schiebel, E. (2018). Brr6 and Brl1 locate to nuclear pore complex assembly sites to promote their biogenesis. J. Cell Biol. jcb.201706024.

Zhang, Y., Luo, C., Zou, K., Xie, Z., Brandman, O., Ouyang, Q., and Li, H. (2012). Single Cell Analysis of Yeast Replicative Aging Using a New Generation of Microfluidic Device. PLoS One.

